# The yeast Art1 arrestin domain contains disordered insertions that regulate its localization and activity

**DOI:** 10.1101/553677

**Authors:** Matthew G. Baile, Evan L. Guiney, Ethan J. Sanford, Jason A. MacGurn, Marcus B. Smolka, Scott D. Emr

## Abstract

The protein composition of the plasma membrane is rapidly remodeled in response to changes in nutrient availability or cellular stress. This occurs, in part, through the selective ubiquitylation and endocytosis of plasma membrane proteins which, in the yeast *Saccharomyces cerevisiae,* is mediated by the HECT E3 ubiquitin ligase Rsp5 and arrestin-related trafficking (ART) adaptors. Here, we provide evidence that an ART family member, Art1, consists of an arrestin fold with extended N- and C-terminal tails, and interspersed with loop insertions. These loop and tail regions, while not strictly required for Art1 function, regulate its activity through two separate mechanisms. One loop mediates Art1 cargo specificity. Other loops are subjected to phosphorylation in a manner dependent on the Pho85 cyclins Clg1 and Pho80. Phosphorylation of the loops controls Art1’s localization to the plasma membrane, which promotes cargo ubiquitylation and endocytosis, demonstrating a mechanism through which Art1 activity is regulated.

## INTRODUCTION

Cells interact with, and respond to, the extracellular environment through the plasma membrane (PM). The PM contains a complex collection of channels, receptors, and transporters and it must be rapidly remodeled to respond to changes in the environment and maintain cellular homeostasis. This occurs through the opposing processes of protein trafficking to the PM and endocytosis. Selective endocytosis occurs via ubiquitin conjugation to a PM protein, which commonly serves as a signal for endocytosis, and lysosomal sorting. The budding yeast *S. cerevisiae* has proved to be a powerful and useful model to understand selective ubiquitin-mediated endocytosis. Numerous nutrient permeases have been shown to be specifically endocytosed and downregulated in response to changes in the extracellular concentration of each permease’s substrate (Gournas et al., 2016; Haguenauer-Tsapis and André, 2004).

In yeast, ubiquitylation of PM proteins is mainly mediated by the HECT-type E3 ubiquitin ligase Rsp5 (Dupré et al., 2004; Lauwers et al., 2010; Léon and Haguenauer-Tsapis, 2009; MacGurn et al., 2012). Rsp5 contains 3 WW domains which specifically recognize PPXY motifs that allow binding and ubiquitylation of substrates (Dunn and Hicke, 2001; Gajewska et al., 2001; Macias et al., 1996). Many Rsp5 substrates, including most PM proteins, do not contain their own PPXY motifs but instead rely on PPXY-containing adaptor proteins to facilitate Rsp5 recruitment and ubiquitylation (Léon and Haguenauer-Tsapis, 2009). Although yeast contain membrane bound Rsp5 adaptor proteins that function on internal membranes (Hettema et al., 2004; Léon et al., 2008; MacDonald et al., 2011; Sardana et al., 2018), most PM proteins are recognized by a family of soluble arrestin-related trafficking (ART) adaptor proteins (Lin et al., 2008; Nikko and Pelham, 2009).

ART proteins belong to the α-arrestin protein family, a conserved protein family that consists of 14 members in yeast and at least 6 in mammals (Alvarez, 2008; Aubry and Klein, 2013). Similar to visual and β-arrestins, α-arrestins contain conserved N- and C-terminal arrestin domains (Aubry et al., 2009). Previous sequence analysis of the ART proteins suggested that most of the ART family members contain an N-terminal arrestin fold and a long C-terminal tail, which contains PPXY motifs (Aubry and Klein, 2013; Becuwe et al., 2012a; Lin et al., 2008).

Each ART protein is able to recruit Rsp5 to its own specific subset of PM cargos in response to specific stimuli, resulting in cargo ubiquitylation and endocytosis (reviewed in (Lauwers et al., 2010). For example, Art1/Ldb19 targets Rsp5 to the methionine permease Mup1 in response to increased extracellular methionine, and the arginine permease Can1 in response to increased extracellular arginine (Ghaddar et al., 2014; Lin et al., 2008); whereas Art3/Aly2 mediates the endocytosis of the aspartic acid/glutamic acid transporter Dip5 in response to increased aspartic acid concentrations (Hatakeyama et al., 2010). Further, multiple ARTs can regulate a single permease, as is the case for the uracil transporter Fur4. Art1, Art2/Ecm21, and Art8/Csr2 all can facilitate Fur4 endocytosis (Nikko and Pelham, 2009). Finally, different stimuli can activate different ARTs, leading to the ubiquitylation the same cargo; the lysine permease Lyp1 is endocytosed in response to increased lysine concentrations via Art1, and in response to cycloheximide via Art2 (Lin et al., 2008). Thus, the ART adaptors recruit Rsp5 to specific cargos in response to specific stimuli, forming a complicated signaling network that tightly and specifically modulates the PM proteome.

It has been proposed that substrate transport facilitates endocytosis both by inducing movement out of specific PM domains to membrane domains where endocytosis is more favored (Busto et al., 2018; Gournas et al., 2018), and by exposing an ART binding site on the permease to allow ubiquitylation to occur (Fujita et al., 2018; Guiney et al., 2016; Keener and Babst, 2013). In addition to regulatory mechanisms involving the permeases, the ARTs themselves are subject to regulation. Most ARTs are ubiquitylated by Rsp5 (Gupta et al., 2007), and for many ARTs this ubiquitylation promotes ART activity (Becuwe et al., 2012b; Hovsepian et al., 2017; Lin et al., 2008). Additionally, Art4/Rod1 can be ubiquitylated at sites separate from its activating ubiquitin which results in proteasomal degradation; a process that is counteracted by deubiquitylating enzymes (Ho et al., 2017). Finally, many ARTs can be regulated by phosphorylation. As examples, Art1 is phosphorylated by Npr1 in response to TORC1 activity (MacGurn et al., 2011), Art4 is phosphorylated by Snf1 (Becuwe et al., 2012b), Art8 is phosphorylated by PKA, and Art6/Aly1 is dephosphorylated by calcineurin (O’Donnell et al., 2013). In every case described so far, phosphorylation inactivates, while dephosphorylation activates, ARTs. The exact mechanism of how phosphorylation regulates the ARTs in currently unclear, although studies with Npr1 suggest that phosphorylation/dephosphorylation cycles regulate the ability of Art1 to associate with the PM (MacGurn et al., 2011). Despite this, much remains to be discovered regarding how ARTs are able to specifically and temporally direct PM cargo ubiquitylation and endocytosis.

Here, we demonstrate that Art1, as well as the other ART proteins in S. *cerevisiae,* likely form an arrestin fold using the full length of their primary sequence, with multiple large insertions between sequences that form the arrestin fold. These “loops” and “tails” are not strictly required for Art1 function, but regulate its activity. Loop 3 confers cargo specificity to Art1, where it is required for Can1 endocytosis but is dispensable for Mup1 and Lyp1 endocytosis. Further, loop 1 and the C-terminal tail, as well as a poorly conserved region within the arrestin fold (termed a “mini-loop”), are phosphorylated in a manner dependent on the Pho85 cyclins Clg1 and Pho80. This phosphorylation inactivates Art1 and prevents cargo endocytosis by reducing the ability of Art1 to associate with the PM.

## RESULTS

### Art1 contains an arrestin domain disrupted by multiple insertions

Sequence analysis and structural predictions suggest that the ART family members typically contain an N-terminal arrestin fold with a PPXY motif-containing C-terminal tail (Aubry and Klein, 2013; Becuwe et al., 2012a; Lin et al., 2008). However, we have previously provided evidence that Mup1 and Can1 interact with Art1 by binding to a motif in the Art1 C-terminal tail, rather than binding within the predicted N-terminal arrestin fold (Guiney et al., 2016). This was unexpected, since the arrestin domain directly binds substrate in a variety of arrestin-domain containing proteins (Aubry and Klein, 2013). Mutating the substrate binding motif (residues R653 and R660 mutated to Asp, termed “*art1*^*2RD*^”) completely ablated Art1 function. Cells expressing *art1^2RD^* were hypersensitive to the toxic arginine analog canavanine (figure 1A). Canavanine is transported into cells via Can1 (Ahmad and Bussey, 1986; Grenson et al., 1966), but toxicity can be mitigated by Can1 endocytosis and vacuolar degradation, and canavanine hypersensitivity occurs when Can1 cannot be endocytosed and degraded, providing a readout of Art1 function (Lin et al., 2008). Additionally, similar to *art1Δ,* cells expressing *art1^2RD^* were unable to grow at 38°C (figure 1A). At elevated temperatures, Art1 is required to endocytose misfolded PM proteins, including Lyp1, preventing membrane permeability and cell death (Zhao et al., 2013). Growth at high temperature is currently the broadest and least sensitive assay for Art1 function. Thus, the inability of *art1^2RD^* to grow at 38°C indicates a severe defect in Art1 function. Further, in cells expressing *art1^2RD^,* methionine-induced Mup1-FLAG degradation was inhibited to a similar extent as in *art1Δ,* whereas Mup1-FLAG degradation was nearly complete when wild type *ART1* was expressed (figure 1B). Thus, R653 and R660 are required for Art1 function. Importantly, Art1^2RD^ fused to mNeonGreen (Art1^2RD^-mNG) was expressed and exhibited wild type-like localization (figure 1C), consistent with a disruption in substrate binding rather than a more general defect in folding or expression.

**Figure 1.**
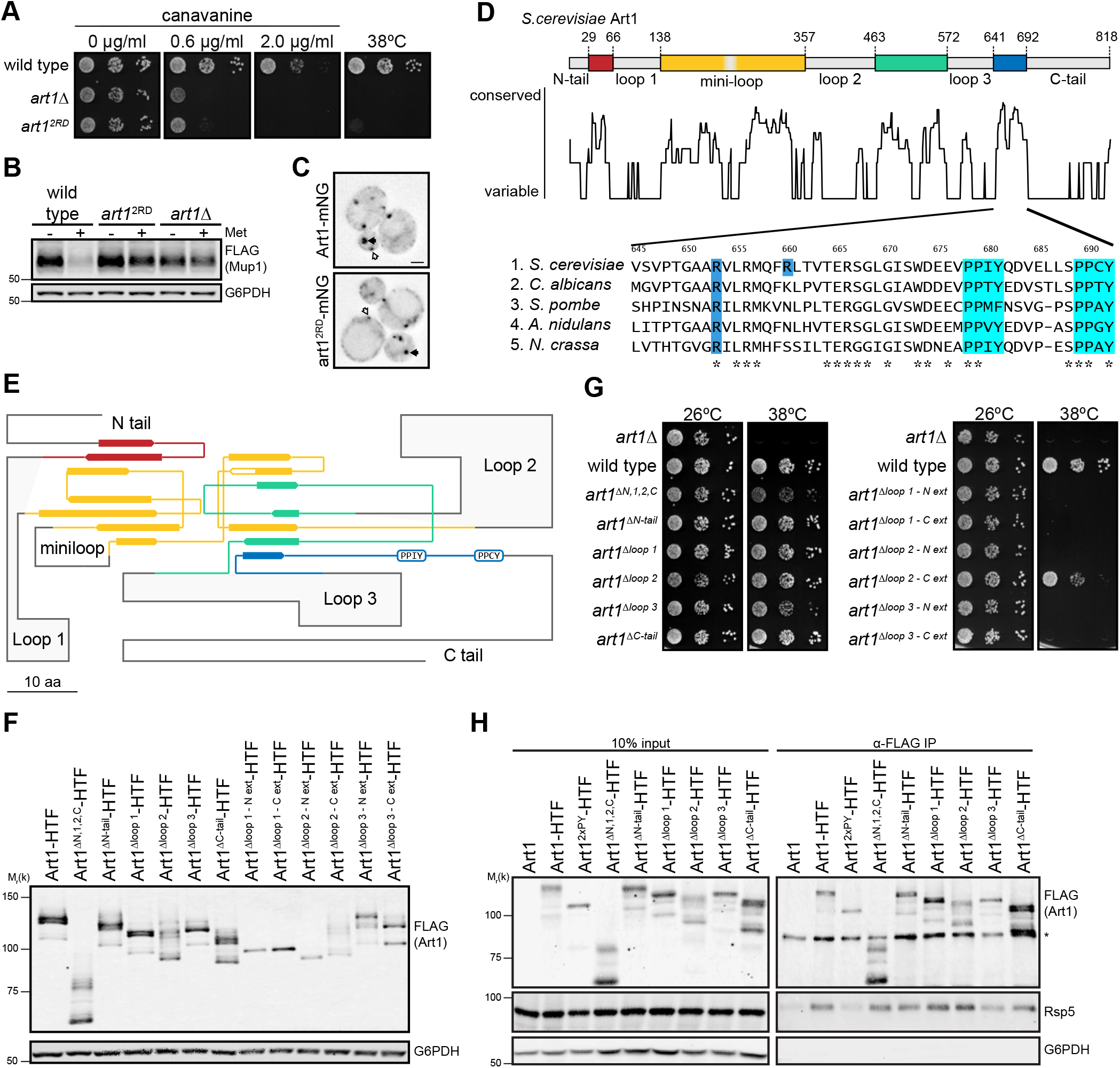
Art1 contains an arrestin domain disrupted by multiple insertions. (A) Serial dilutions of *art1Δ* yeast, or *art1Δ* yeast expressing ART1 (wild type) or *art1^2RD^* (*art1*^R653D,R660D^), spotted onto synthetic media containing the indicated concentration of canavanine, and incubated at 26°C or 38°C. (B) The yeast strains in A containing Mup1-FLAG were treated with 20 μg/mL Met for 60 min before protein extraction. Cell lysates separated by SDS-PAGE and immunoblotted with the indicated antibodies. (C) Localization of Art1-mNG and art1^2RD^-mNG in minimal media. White arrows indicate PM localization, black arrows indicate Golgi localization. Scale bar = 1 μm. (D) (top) Art1 schematic. Non-conserved loop regions are labeled and shown in grey. Conserved regions predicted to form an arrestin fold are colored. (middle) relative conservation of *S.cerevisiae* Art1 calculated with rate4site. (bottom) Sequence alignment of a well conserved C-terminal region of Art1. Blue highlights indicate the predicted substrate binding residues, R653 and R660. Cyan highlights indicate the Rsp5 binding PPXY motifs. (E) Secondary structure diagram of Art1 showing the organization of the β-sheets relative to the loops and tails (F) Steady state expression analysis of Art1 tail, loop, and loop extension mutants. Cell lysates separated by SDS-PAGE and immunoblotted with the indicated antibodies. (G) Serial dilutions of the strains in G spotted on synthetic media, incubated at 26°C or 38°C. (H) α-FLAG immunoprecipitation of HTF-tagged Art1 loop and tail mutants. * indicates a non-specific band detected by the α-FLAG antibody.

To further investigate this C-terminal substrate binding region, we analyzed the conservation of this region among all the fungal Art1 homologs. The region surrounding R653 and R660 is well conserved among the Art1 homologs examined, suggesting that they all bind their substrates using a similar motif. Strikingly, applying this analysis to the entire sequence revealed that Art1 (and its closely related homologs) contained multiple large, variable insertions (herein referred to as “loops” and “tails”) between regions that were well conserved. Shorter, more distantly related Art1 orthologs retained only the well conserved regions, while the loops and tails were absent (figure 1D, figure 1 – supplement 4). As an example, the distantly related Any1/Arn1 from S. *pombe* contains only a short N-terminal tail, and no loops (figure 1 – supplement 2). Structural modeling predicts that Any1 still forms an arrestin fold (figure 1F), suggesting that the inserted loops and tails in Art1 are not part of the arrestin domains (figure 1 – supplement 2). In the new structural model, loop 1 is inserted between β-strand 2 and 3 in the N-lobe, loop 2 is positioned between the 3^rd^ and 4^th^ β-strand of the C-lobe, and loop 3 occurs before the final β-strand in the C-lobe (figure 1E). There is also a shorter non-conserved region, termed a “mini-loop,” which also maps to a turn between two β-strands. Interestingly, the regions where the loops are predicted to be inserted occur in turns between β-strands or coils, allowing them to extend away from the arrestin domain, and therefore are not expected to disrupt the core arrestin fold. Thus, Art1 may form its arrestin fold using the conserved “core” regions with interspersed loops, rather than forming an arrestin fold with its N-terminal half. Further supporting this model, the variable loops and tails are predicted to be disordered, unlike the conserved core regions, suggesting that these regions do not tightly fold into a structured domain (figure 1 – supplement 1).

To test this hypothesis, the N- and C-terminal tails, and each loop, were removed individually and tested for function. Steady state expression of each loop mutant was similar to wild type Art1 (figure 1F) and each mutant could grow at elevated temperature (figure 1G), demonstrating that the loop and tail deletions did not completely ablate Art1 function. When the N-terminus, loop 1, loop 2, and the C-terminus deletions were combined (termed “*art1*^ΔN,1,2,C^”), cells exhibited a reduced ability to grow at elevated temperature, but not to the extent of *art1Δ,* demonstrating that many of the loops and tails contribute to, but are not strictly required for, Art1 function. Coimmunoprecipitation analysis showed that each loop and tail mutant was able to interact with Rsp5, unlike the PPXY mutant *(art1^2xPY^*) which was unable to bind Rsp5 (figure 1H). Thus, removing these relatively large regions of Art1 does not prevent folding so severely that it is unable to form its normal protein-protein interactions. Consistently, each loop and tail mutant was ubiquitylated (figure 1F), which requires binding to Rsp5 (Lin et al., 2008). Although attempts to model the structure of full length Art1 were unsuccessful, modeling the minimal Art1^ΔN,1,2,C^ mutant predicts a full arrestin fold (figure 1 – supplement 3). Additionally, extending each loop deletion 10-15 additional amino acids into the predicted core arrestin fold completely abolished Art1 activity (figure 1G), except for the loop 2 C-terminal extension, further supporting the identification of the predicted loops (with the exception of the precise loop 2 C-terminal boundary). Both loop 1 extension mutants and the loop 2 N-terminal extension mutant were not ubiquitylated (figure 1F), suggesting that protein folding was disrupted to the extent that Rsp5 binding was affected, despite retaining the PPXY motifs. However, both loop 3 extension mutants were ubiquitylated, signifying that Art1 function was abolished through a mechanism distinct from decreased Rsp5 binding. Taken together, these data support a model where the arrestin fold of Art1 consists of multiple conserved regions that are disrupted by long insertions.

Interestingly, analysis of the other *S. cerevisiae* ART family members indicates a similar theme, where rather than containing one contiguous arrestin fold in their N-terminus, they instead contain poorly conserved insertions between conserved regions that make up the arrestin domain. Strikingly, for each ART the size and position of the insertions differs, suggesting that the arrestin domain can tolerate insertions at many places while still maintaining the ability to properly fold (figure 1 – supplement 4–7, extended data file 1).

### Art1 loops regulate cargo endocytosis

Since individually deleting the Art1 loops and tails did not result in a measurable defect in Art1-mediated PM quality control, but the combined *art1^ΔN,1,2,C^* mutant did show a decrease in function (figure 1G), we hypothesized that the loops and tails could be involved in regulating Art1. We further examined the effect of the loop and tail mutants on two Art1 cargos, Can1 and Mup1. When grown on canavanine, the strains expressing *art1^ΔN,1,2,C^* and *art1^Δloop3^* were canavanine hypersensitive (figure 2A) implying a Can1 endocytosis defect, while the remaining mutants exhibited wild type-like or slightly impaired canavanine growth. This was consistent with Can1-GFP degradation after addition of its substrate, Arg (figure 2B, figure 2 – supplement 1). Can1-GFP degradation occurs after treatment with Arg in a dose-dependent manner, thus higher Arg concentrations applied for the same amount of time cause more Can1-GFP degradation in wild type cells. Cells expressing *art1^ΔN,2,C^* exhibited less Can1-GFP degradation at higher Arg concentrations (20 and 100 μg/mL) compared to wild type *ART1*, whereas *art1^Δloop 3^* expression caused a more complete block in Can1-GFP degradation that was indistinguishable from that observed in *art1Δ.* The remaining loop deletion mutants showed a degradation profile similar to that of wild type *ART1.*

**Figure 2.**
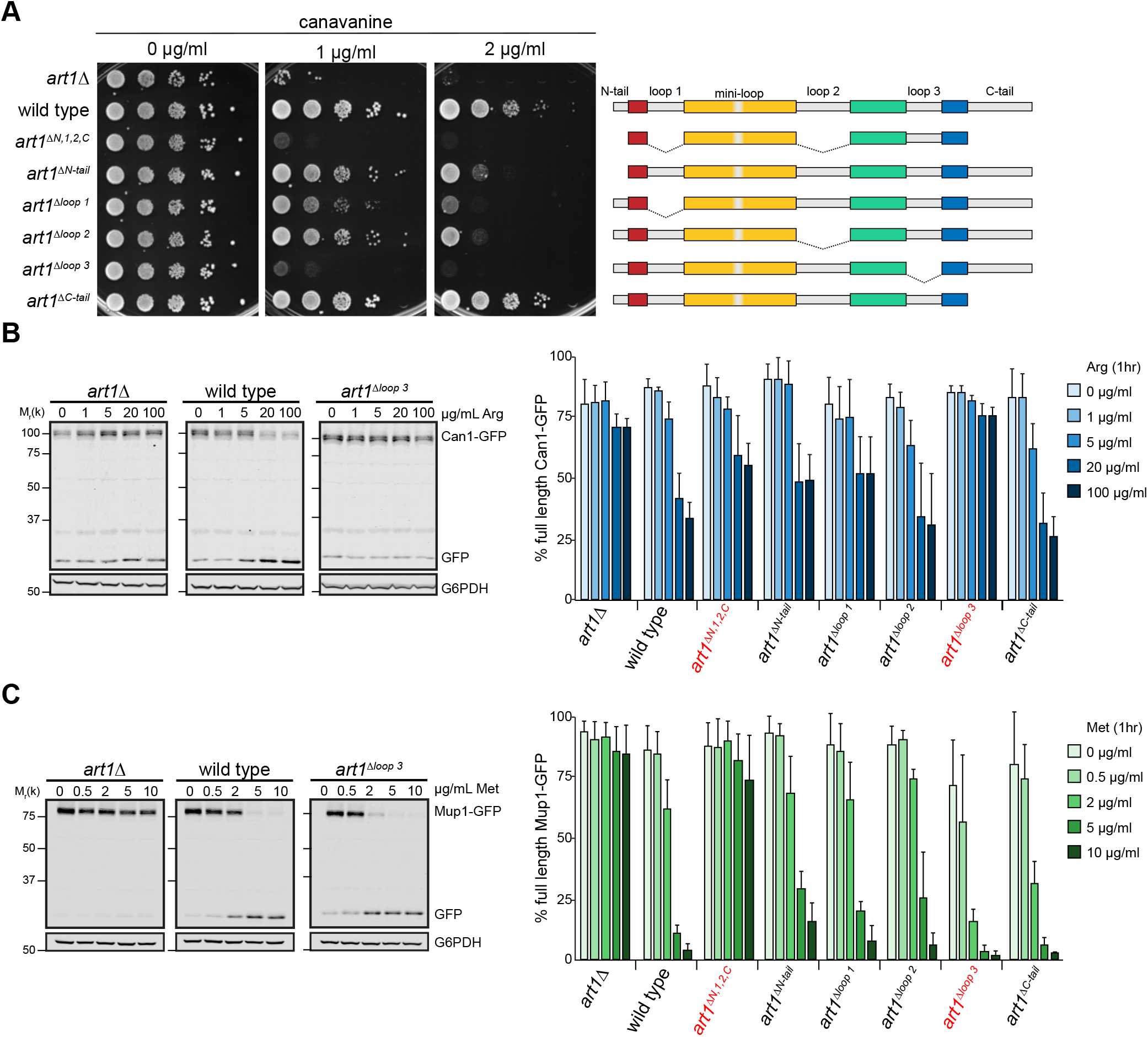
Art1 loop 3 dictates substrate specificity. (A) Serial dilutions of the Art1 tail and loop mutant strains spotted on synthetic media containing the indicated concentration of canavanine, incubated at 26°C. (B) Immunoblot analysis of Can1-GFP endocytosis inducted with the indicated concentration of Arg for 60 min in *art1* Δ cells or *art1Δ* cells expressing *ART1* or *art1Δ^¤p3^* (left). Band intensities from three biological replicates were quantified and expressed as the mean % full length Can1-GFP remaining (right). Bars indicate 95% confidence intervals. (C) Immunoblot analysis of Mup1-GFP. As in B, except Met was used to induce endocytosis.

Growth on thialysine, a toxic analog of Lys that is a substrate of the Lyp1 transporter, was consistent with the growth on canavanine. Cells expressing *art1^ΔN,1,2,C^* and *art1^Δloop 3^* were the most thialysine hypersensitive, with less severe defects exhibited by cells expressing *art1^ΔN-tal^* and *art1^Δloop 1^* (figure 2 – supplement 2).

For most loops and tails mutants, Met-induced Mup1-GFP degradation followed the same approximate pattern as Can1-GFP degradation. Most mutants displayed a Mup1-GFP degradation profile similar to wild type, but *art1^ΔN,1,2,C^* caused a strong block in degradation (figure 2C, figure 2 – supplement 3). Together with the Can1-GFP data, this demonstrates that while deleting the individual tails or loops had little effect, combining the deletions resulted in a strong block of cargo-specific endocytosis. Notably, *art1^ΔC-tail^* caused more complete Mup1-GFP degradation at lower Met concentrations, which was observed to a much lesser extent for Can1-GFP endocytosis.

Interestingly, and in contrast to Can1-GFP degradation, cells expressing *art1^Δloop3^* exhibited enhanced Mup1-GFP degradation at lower concentrations compared to *ART1*. Combined with the hypersensitivity to thialysine (figure 2 – supplement 2) and the ability to grow at elevated temperature (figure 1G), these data support a model where loop 3 is required for Can1 and Lyp1 endocytosis but is dispensable for Mup1 endocytosis and PM quality control. Therefore, loop 3 contributes to Art1 function by dictating substrate specificity.

### Art1 loops regulate its localization

To investigate the role the remaining loops and tails may be playing in Art1 regulation, we analyzed Art1 localization in these mutants (figure 3A). Art1 is predominately localized to the Golgi but upon nutrient stimulation translocates to the PM (Lee et al., 2019; Lin et al., 2008; MacGurn et al., 2011; Martínez-Márquez and Duncan, 2018), presumably allowing Art1 to bind to its cargo, resulting in ubiquitylation and endocytosis. One such stimulus for Art1 PM association is shifting cells from minimal synthetic media (SCD) to rich media (YPD). Upon shifting to rich media, the intensity of Art1-mNG at the PM increases (figure 3B, figure 3 – supplement 1), with a concurrent decrease in intensity at the Golgi (figure 3 C, figure 3 – supplement 1). By measuring the Art1-mNG intensity at both the PM and Golgi, we determined the localization of each Art1 loop mutant in minimal media and after 60 min in rich media (figure 3).

**Figure 3.**
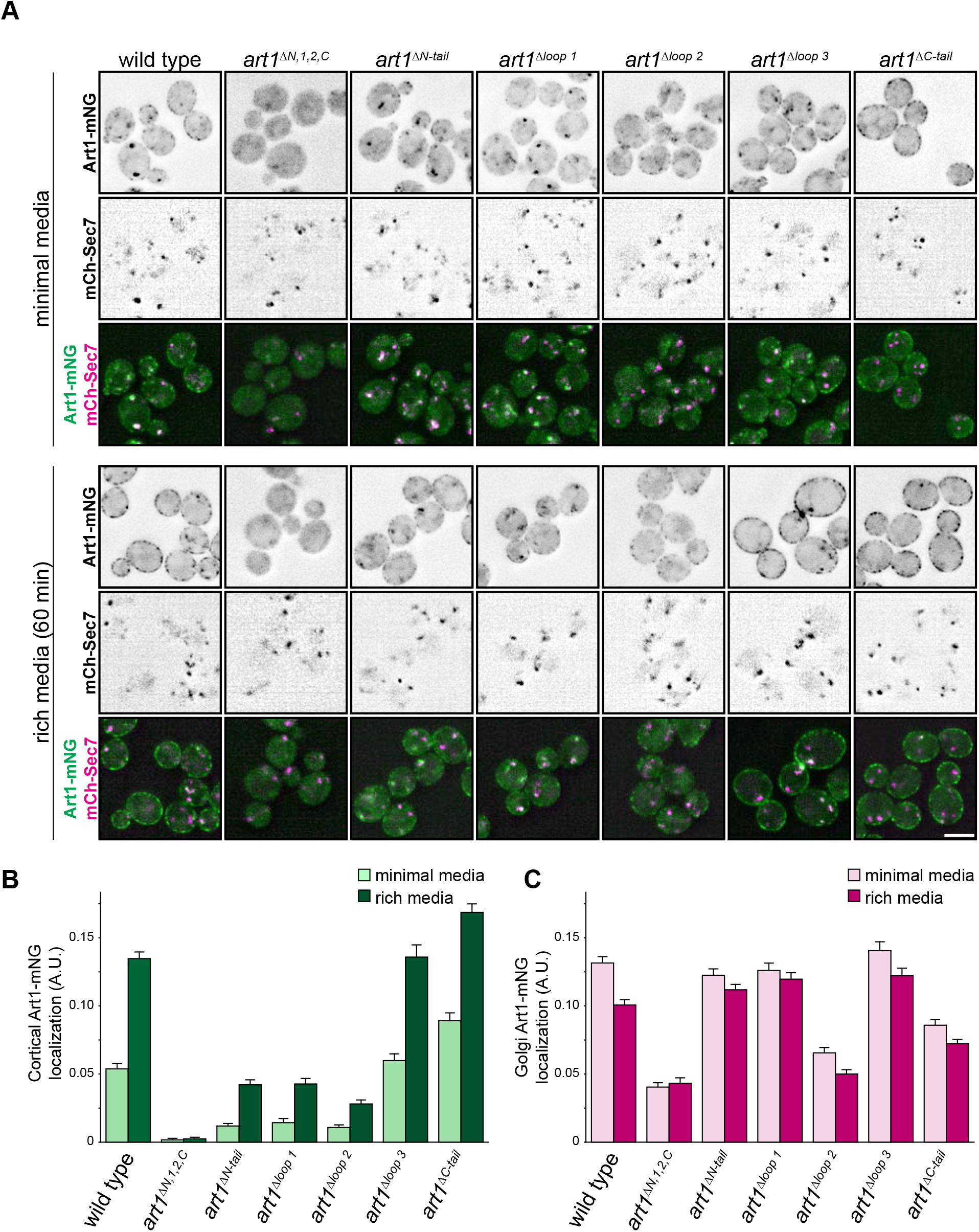
Localization of Art1 loop and tail mutants. (A) Localization of the indicated Art1-mNG loop and tail mutants in minimal media and after shifting to rich media for 60 min. Bar = 2 μm. (B) Quantification of PM localization of the indicated Art1 loop and tail mutants. (C) Quantification of Golgi localization of the indicated Art1 loop and tail mutants. Bars indicate 95% confidence intervals

The Golgi localization of Art1^Δloop3^-mNG in minimal media, and its ability to associate with the PM after treatment with rich media, was similar to wild type Art1-mNG, demonstrating that its cargo-specific regulation occurs through a mechanism distinct from altered subcellular localization.

Art1^ΔC-tail^-mNG, even in minimal media, shows more PM localization than wild type, and is accompanied by a decrease in Golgi localization, consistent with its ability to cause cargo endocytosis at lower substrate concentrations (figure 2). Thus, the Art1 C-tail promotes its Golgi localization, as its absence leads to increased Art1 on the PM.

Art1^ΔN,1,2,C^-mNG is completely unable to localize to the PM (figure 3A and B). This decreased ability to localize to the PM, and thus bind PM cargo, is the likely mechanism explaining the defect in cargo endocytosis (figure 2). Art1^ΔN,1,2,C^-mNG also has a weaker association with the Golgi (figure 3C), suggesting that it is deficient for all membrane association, not just the PM. Notably, the combination of N-terminal tail, loop 1, and loop 2 were able to overcome the PM localization caused by deleting the C-terminal tail. Interestingly, the individual N-terminal tail, loop 1, and loop 2 mutants each retain some ability to associate with the PM after stimulation with rich media. Although the extent of total PM signal is reduced compared to wild type, these mutants can still mediate wild type-like cargo endocytosis (figure 2). Therefore, the major localization defect observed for Art1^ΔN,1,2,C^-mNG is due to additive effects of each individual loop deletion.

### Art1 regulation by Pho85

The effect of the large deletions of Art1 loops and tails on Art1 localization are consistent with the loops and tails participating in Art1 regulation. Previous work has demonstrated that Art1 is regulated by Npr1-mediated phosphorylation (MacGurn et al., 2011), on residues in loop 1 (figure 1D). However, not all of the identified residues were dependent on Npr1, suggesting other kinases may regulate Art1 function. Thus, we tested various kinase knockout mutants for growth on canavanine and found that *pho85Δ* exhibited canavanine resistance (figure 4A). Pho85, the yeast homolog of Cdk5 (Carroll and O’Shea, 2002), is a cyclin-dependent kinase, and 10 cyclins have been identified to direct Pho85 to its substrates (Measday et al., 1997; Moffat et al., 2000). Of the individual cyclin mutants, *clg1Δ* was canavanine resistant, and *pho80Δ* was canavanine hypersensitive (figure 4A), whereas the remaining cyclin mutants grew like wild type on canavanine (figure 4 – supplement 1), suggesting that Clg1 inhibits Art1 and Pho80 activates it. Interestingly, because both Clg1 and Pho80 are dependent on Pho85 but exhibit opposite canavanine phenotypes, the canavanine sensitivity phenotype of *pho85*Δ was intermediate to either *clg1Δ* or *pho80Δ* (figure 4A). When the cyclin mutants were tested for growth on thialysine, *clg1Δ* and *pho80Δ* recapitulated their canavanine phenotypes, with *clg1Δ* being resistant and *pho80Δ* being hypersensitive (figure 4 – supplement 2). Importantly, Art1 expression was unaffected when *PHO85, PHO80,* or *CLG1* were deleted (figure 4B). Additionally, *CLG1* overexpression caused cells to become canavanine hypersensitive (figure 4C), consistent with Clg1 inactivating Art1.

**Figure 4.**
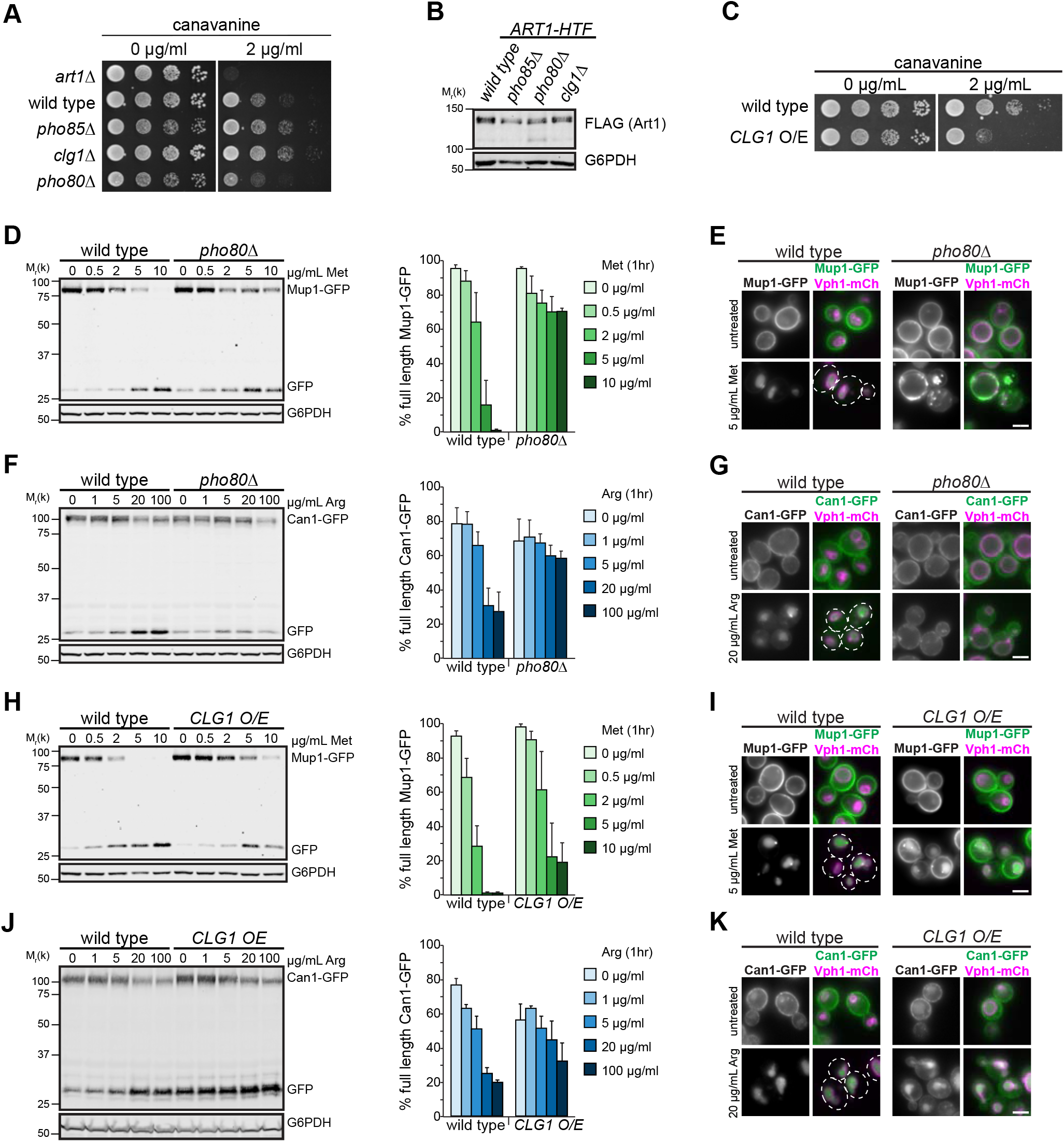
Pho85, and the cyclins Pho80 and Clg1, regulate Art1 activity. (A) Serial dilutions of the indicated null mutants on spotted on synthetic media containing the indicated concentration of canavanine and incubated at 26°C. (B) Steady state expression of Art1 in the indicated null mutants analyzed by immunoblot. (C) Serial dilutions of wild type or cells overexpressing *CLG1* spotted on synthetic media containing the indicated concentration of canavanine and incubated at 26°C. (D) Immunoblot analysis of Mup1-GFP endocytosis induced with the indicated concentration of Met for 60 min in wild type or *pho80Δ* yeast. Band intensities from three biological replicates were quantified and expressed as the mean % full length Mup1-GFP remaining. Bars indicate 95% confidence intervals. (E) Fluorescence microscopy of wild type or *pho80Δ* cells expressing Mup1-GFP, with and without inducing endocytosis by treating with 5 μg/mL Met for 60 min. (F) As in D, except analyzing Can1-GFP endocytosis induced with the indicated concentration of Arg for 60 min. (G) As in E, except analyzing Can1-GFP localization after inducing endocytosis with 20 μg/mL Arg for 60 min. (H) As in D, except comparing wild type yeast to cells overexpressing *CLG1.* (I) As in E, except comparing wild type yeast to cells overexpressing *CLG1.* (J) As in F, except comparing wild type yeast to cells overexpressing *CLG1.* (K). As in G, except comparing wild type yeast to cells overexpressing *CLG1.* Bar = 2 μm

In the absence of *PHO80,* Art1-mediated endocytosis is impaired. While Mup1-GFP was almost completely degraded after 60 min of treatment with 5 or 10 μg/ml Met in wild type cells, in *pho80Δ* only ~40% of Mup1-GFP was degraded, even at higher Met concentrations (figure 4D). Consistently, Mup1-GFP was completely sorted to the vacuole in wild type cells after Met treatment, whereas in *pho80Δ* a portion of Mup1-GFP was found either on internal puncta, presumed to be endosomes, or was retained on the PM (figure 4E). Importantly, this demonstrates that the defect in Mup1-GFP degradation was due to an endocytosis and vacuole sorting defect rather than a defect in vacuolar proteolysis. Analysis of Can1-GFP endocytosis revealed the same trend. Even at higher concentrations of Arg, Can1-GFP degradation was impaired in *pho80Δ* compared to wild type cells (figure 4F), and Can1-GFP was mostly retained on the PM in *pho80Δ* cells after treatment with Arg (figure 4G), although some signal was observed on endosomes and (to a much lesser extent) in the vacuole lumen. Taken together with the hypersensitivity of *pho80Δ* cells to canavanine (figure 4A), these data indicate that Pho80 positively regulates Art1, resulting in an Art1-mediated endocytosis defect in *pho80Δ.*

In the absence of *CLG1,* there was a slight increase in Mup1-GFP degradation at intermediate Met concentrations (figure 4 – supplement 3), which was undetectable by microscopy, even at shorter timepoints (figure 4 – supplement 4). Interestingly, there was no increase in constitutive, non-Met induced endocytosis, as might be expected. Can1-GFP degradation and vacuolar sorting in *clg1Δ* was indistinguishable from wild type (figure 4 – supplement 5 and 6). However, *CLG1* overexpression, resulted in a Mup1-GFP degradation defect (figure 4H) and a delay in vacuolar sorting (figure 4I). Similar effects were observed for Can1-GFP degradation and endocytosis (figure 4J and K). Although a striking increase in cargo endocytosis was not observed in *clg1Δ,* despite *clg1Δ* cells exhibiting canavanine resistance, this may reflect the inherent differences in the sensitivity of these assays. However, *CLG1* overexpression did cause an endocytosis defect, consistent with Clg1 activity inactivating Art1.

### Pho80 and Clg1 affect Art1 localization

Since Pho80 and Clg1 activate and inactivate Art1, respectively, we next sought to understand the mechanism(s) by which they modulated Art1 function. Thus, we examined Art1 localization in the mutants. In *pho80Δ* cells grown in minimal media, Art1-mNG remained mostly localized at the Golgi, similar to wild type. However, after an acute treatment with rich media, Art1-mNG was unable to efficiently translocate to the PM in the same way as in wild type cells (figure 5A), suggesting that the Art1-mediated endocytosis defect observed in *pho80Δ* cells is the result of Art1 failing to associate with the PM.

**Figure 5.**
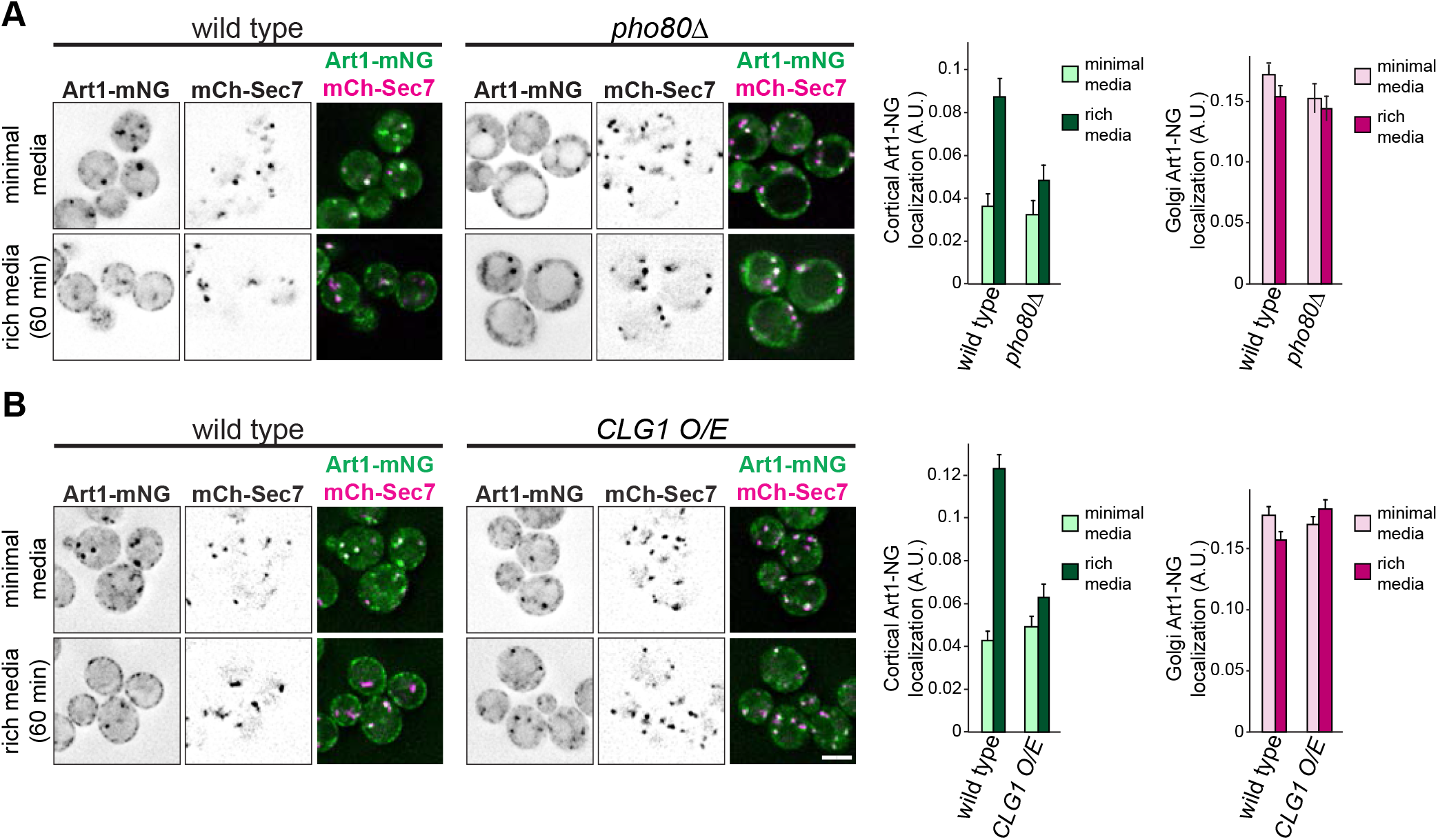
Pho80 and Clg1 affect Art1 localization. (A) Localization of Art1-mNG in wild type and *pho80Δ* yeast in minimal media and after shifting to rich media for 60 min. (right) quantification of Art1-mNG PM and Golgi localization. Bars indicate 95% confidence intervals. (B) As in A, except comparing Art1-mNG in wild type and cells overexpressing *CLG1.* (right) quantification of Art1-mNG PM and Golgi localization. Bars indicate 95% confidence intervals. Bar = 2 μm.

Art1-mNG localization in *clg1Δ* mimicked its localization in wild type cells (figure 5 – supplement 1), consistent with the cargo endocytosis phenotypes (figure 4 – supplements 3–6). However, *CLG1* overexpression caused a strong defect in Art1-mNG PM localization in rich media (figure 5B). Therefore, Clg1 decreases Art1 activity by preventing or decreasing Art1 PM localization. Thus, both Clg1 and Pho80 regulate Art1 in a manner similar to some of the loops and tails mutants: by modulating its PM association.

### Phosphorylation on loops and tails regulate Art1

Considering that loop 1 in Art1 is a known target for Npr1-mediated phosphorylation (MacGurn et al., 2011), and that Clg1 and Pho80 can regulate Art1 localization reminiscent of the loop and tail mutants, we hypothesized that the loops and/or tails are phosphorylated in a Clg1-and Pho80-dependent manner. Thus, we identified the Clg1- and Pho80-mediated phosphorylated residues on Art1 using SILAC mass spectrometry (figure 6A, extended data file 2).

**Figure 6.**
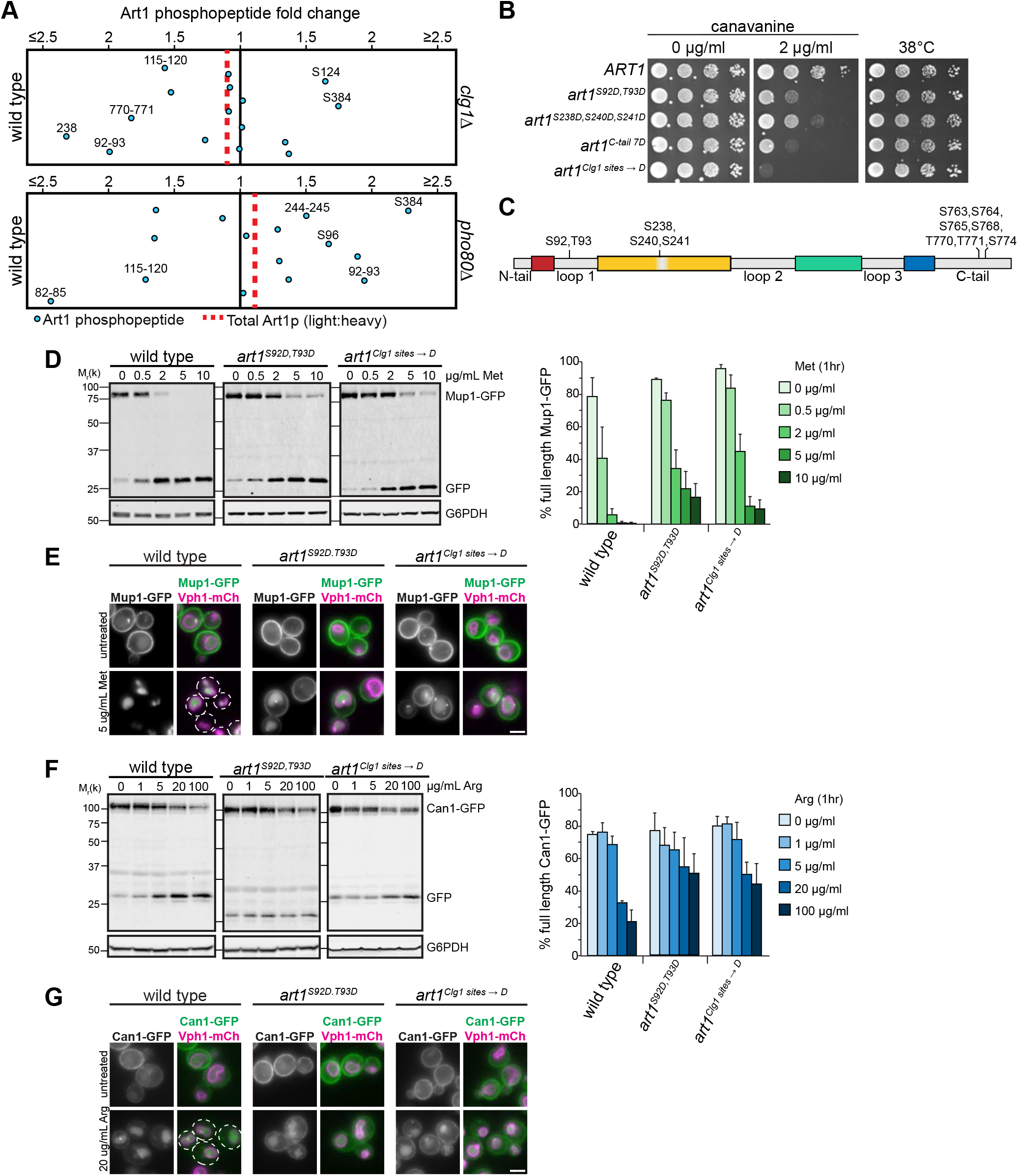
Phosphorylation on loops and tails regulate Art1. (A) QMAPS depicting the fold change in phosphopeptide abundance of Art1 in (top) wild type vs *clg1Δ* or (bottom) wild type vs *pho80Δ.* (B) Serial dilutions of cells expressing the indicated Art1 mutants spotted on synthetic media containing the indicated concentration of canavanine and incubated at 26°C or 38°C. (C) Schematic of Art1 showing the location of the phosphorylated residues that when mutated affect Art1 function. (D) Immunoblot analysis of Mup1-GFP endocytosis induced with the indicated concentration of Met for 60 min in cells expressing wild type *ART1, art1*^S92D,T93D^, or *art1^Clg1 sites→D^*. Band intensities from three biological replicates were quantified and expressed as the mean % full length Mup1-GFP remaining. Bars indicate 95% confidence intervals. (E) Fluorescence microscopy of Mup1-GFP, with and without inducing endocytosis by treating with 5 μg/mL Met for 60 min, in cells expressing wild type *ART1, art1^S92DT93D^,* or *art1^Clg1 sites~ D^*. (F) As in D, except analyzing Can1-GFP endocytosis induced with the indicated concentration of Arg for 60 min. (G) As in E, except analyzing Can1-GFP localization after inducing endocytosis with 20 μg/mL Arg for 60 min. Bar = 2 μm.

In *clg1Δ*, Art1 phosphorylation was reduced at S92/T93, S238, and T770/T771 (figure 6A), suggesting that Clg1 was responsible for these phosphorylation events. Residues S92/T93 are contained within loop 1. S238 is in the middle of a predicted arrestin core region, but interestingly, falls within the poorly conserved “mini-loop” rather than the arrestin fold. T770/T771 are in the C-terminal tail of Art1. Notably, phosphorylation was also modestly decreased on 82-STTS-85 (extended data file 2), residues previously identified as Npr1 targets (MacGurn et al., 2011). As previously hypothesized, there may be cross-talk between Npr1 and other kinases; in this case, Clg1.

Mutating S92 and T93 to the phosphorylation mimetic Asp resulted in canavanine hypersensitivity (figure 6B), consistent with the *CLG1* overexpression phenotype. Similarly, mutating S238 (in combination with the nearby S240 and S241) also caused canavanine hypersensitivity, although less so that the S92D/T93D mutant.

Mutating T770/T771 did not affect growth on canavanine (figure 6 – supplement 1). However, additionally mutating five adjacent Ser residues (all within 15 amino acids of T770/T771) to Asp (termed *“art1^C-tail 7D^”)* caused canavanine hypersensitivity (figure 6B). Thus, similar to Npr1-mediated phosphorylation of loop 1 on Art1 (MacGurn et al., 2011) and the N-terminus of Art8 (Hovsepian et al., 2017), on the Art1 C-tail it is necessary to mutate phosphorylated residues identified by mass spectrometry as well as adjacent residues before a phenotype is revealed, presumably due to inexact phosphorylation of Art1 by its kinase. Notably, mutations to the phosphorylation deficient Ala did not result in canavanine resistance (figure 6 – supplement 2).

Hence, three distinct and distant regions of Art1 are phosphorylated in a Clg1-dependent manner: loop 1, a “mini-loop” in core region 2, and the C-terminal tail (figure 6C). While these data cannot determine if all three of these regions are regulated simultaneously or if they represent individually regulated sub-populations of Art1, combining the phosphorylation mimetic mutants of all three regions (termed *“art1^Clg1 sites →D^*”) resulted in an even greater canavanine sensitivity than the individual mutants (figure 6B), demonstrating additive effects of phosphorylation at these regions of Art1. Notably, the steady state expression levels of each mutant is similar to wild type (figure 6 – supplement 3). Importantly, while canavanine sensitivity is affected by these phosphorylation site mutants, the ability of Art1 to facilitate growth at elevated temperature remains intact, suggesting that while Clg1-medated phosphorylation can regulate substrate mediated endocytosis of Can1, it does not block Art1-mediated PM quality control.

When Art1 phosphorylation in *pho80Δ* was analyzed, only phosphorylation at residues 82-STTS-85 was significantly decreased (figure 6A). These sites have previously been shown to be phosphorylated by Npr1. Perplexingly, dephosphorylation at these sites results in canavanine resistance and Art1 activation (MacGurn et al., 2011), but *pho80Δ* cells are canavanine hypersensitive (figure 4A). Intriguingly, multiple residues become hyperphosphorylated in *pho80Δ,* including S92/T93. The phosphorylation mimetic Asp mutants at S92/T93 resulted in Art1 inactivation (figure 6B), which is consistent with the *pho80Δ* phenotype (figure 4A) and suggests that phosphorylation at S92/T93 can inactivate Art1 even when the Npr1 sites are dephosphorylated. Hence, Pho80 activity regulates Art1, but it does so indirectly, likely by activating another kinase or by inactivating a phosphatase that directly interacts with Art1. S384 was also hyperphosphorylated in *pho80Δ* (figure 6A). However, neither mutating S384 to Ala or Asp resulted in any change in canavanine sensitivity. Combining S384 mutants with S92/T93 mutants perfectly mimicked the S92/T93 mutant phenotypes (figure 6 – supplement 4), demonstrating that, while S384 is phosphorylated in a Pho80-dependent manner, it has no effect on Can1 endocytosis.

Consistent with the canavanine growth phenotypes, Mup1-GFP degradation was impaired in cells expressing *art1^S92D,T93D^* and *art1^Clg1 sıtes *D^* (figure 6D), and a portion of Mup1 – GFP remained on the PM or, to a lesser extent, at endosomes (figure 6E). Also consistent with the growth on canavanine, Mup1-GFP degradation was like wild type in cells expressing *art1^S92A,T93A^* and *art1^Clg1 sites → A^*, and no constitutive endocytosis was observed in the absence of Met treatment (figure 6 – supplement 5 and 6). Further, cells expressing *art1^S238D,S240D,S241D^* or *art1^C-tail 7D^* showed more modest defects in Mup1-GFP degradation (figure 6 – supplement 5) and vacuole sorting (figure 6 – supplement 6). Similar effects were observed with Can1-GFP endocytosis and degradation when *art1* phosphorylation mutants were expressed (figure 6F and G, figure 6 – supplement 7 and 8). Thus, Clg1- and Pho80-dependent phosphorylation of Art1, although it is indirect, regulates cargo endocytosis.

Clg1 and Pho80 regulate Art1 via its subcellular localization (figure 5), thus we tested if Art1-mNG localization was altered when the Clg1- and Pho80-dependent phosphorylation sites were mutated. Cells expressing *art1^S92D,T93D^* and *art1^Clg1 sites →D^* demonstrated less PM localization in minimal media, and both had defects in the ability to associate with the PM after shifting to rich media (figure 7). While *art1^Clg1 sites →D^* was able to increase its PM localization after shifting to rich media, the maximal level of PM localization was approximately the same as wild type cells in minimal media. Thus, the PM association of *ART1* phosphorylation mimetic mutants is defective. Consistent with all of the Ala mutants at Art1 phosphorylation sites phenocopying wild type, no Ala mutant resulted in constitutive PM localization, and all were able to associate with the PM after acute shift to rich media (figure 7 – supplement 1). Taken together, these data suggest a mechanism by which Clg1- and Pho80-dependent phosphorylation regulates Art1 activity via its ability to associate with the PM.

**Figure 7.**
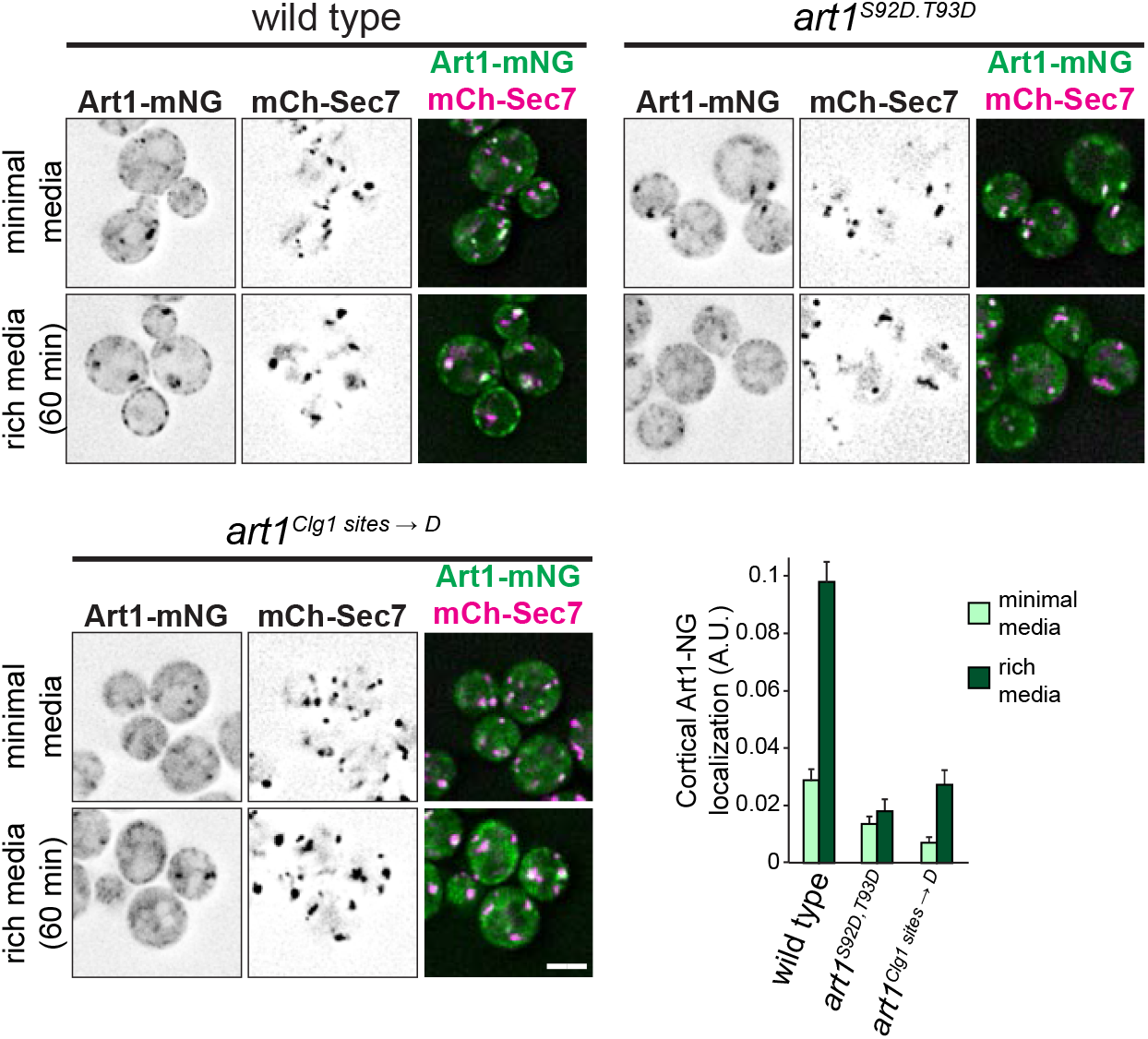
Phosphorylation affects Art1 localization. Localization of wild type Art1-mNG, art1^S92D,T93D^-mNG, or art1^Clg1 sites → D^-mNG in minimal media and after shifting to rich media for 60 min. Bar = 2 μm. (lower right) quantification of Art1 cortical localization. Bars indicate 95% confidence intervals.

## DISCUSSION

This work provides new insight into the mechanisms regulating Art1-mediated endocytosis. Specifically, we show that (1) Art1 forms an arrestin fold using the entire length of its sequence with interspersed and poorly conserved “loop” and “tail” insertions, rather than just its N-terminus, as was previously predicted (2) that the Art1 loops and tails regulate Art1 function via at least two separate mechanisms, by specifying substrate recognition and by controlling Art1 PM translocation, and (3) that phosphorylation on the loops and tail regions regulates Art1 localization.

### A new model for the Art1 arrestin fold

Our sequence analysis of Art1 fungal homologs suggests that Art1 contains multiple large loops, and N- and C-terminal tails, between conserved, arrestin-fold forming regions (figure 1D). This is a striking deviation from previous structural predictions where only the N-terminal half of Art1 formed an arrestin domain (Aubry and Klein, 2013; Becuwe et al., 2012a; Lin et al., 2008). However, deleting the loops and tails retained the ability of Art1 to support growth at elevated temperature (figure 1G), and retains Rsp5 binding (figure 1H), providing evidence that the loop and tail regions are separate from the arrestin fold.

Structural modeling predicts that the loops are positioned in turns between β-strands or coils (figure 1E, figure 1 – supplements 2 and 3), potentially explaining both the lack of conservation of the loops, and how the loops can be removed without completely abolishing Art1 function. Importantly, most mutants where the loop deletions were extended into the conserved sequences of Art1 completely abolished Art1 function (figure 1G), providing additional support for the new structural prediction. This new model places the predicted substrate binding motif on Art1 (figure 1, Guiney et al., 2016) on the arrestin domain, rather than on a distal C-terminal region, consistent with other examples of arrestin domains binding to substrates (Aubry et al., 2009).

Conservation analysis of other ART family members indicated that they, like Art1, also contain non-conserved loops and tails. Strikingly, these insertions occur at different positions relative to the conserved arrestin fold for each ART cluster (Art2/Art8, Art3/Art6, and Art4/Art5/Art7; figure 1 – supplement 4–7). This suggests that large insertions can be tolerated at multiple places within the core arrestin domain without disrupting folding. Considering the regulatory role of the tails and loops in Art1, this highlights potential regulatory regions in other ART family members as well. Interestingly, Art8 is regulated by PKA through phosphorylation on its N-terminus (Hovsepian et al., 2017), which is poorly conserved among fungi (figure 1 – supplement 5); and Art3 is regulated by calcineurin (O’Donnell et al., 2013) by dephosphorylating residues that map to other poorly conserved regions (figure 1 – supplement 6). Thus, it is possible that phosphorylation of poorly conserved loop regions interspersed between conserved arrestin fold regions is a recurring mode of regulation in the ART family.

### Art1 loops and tails as regulatory elements

Art1 loops and tails contribute to its function through two distinct mechanisms: by dictating substrate specificity, and by regulating its localization. Loop 3 is required for Can1 degradation, but not for Mup1 (figure 2). In fact, Mup1-GFP degradation occurs more completely at lower Met concentrations when loop 3 is deleted than with wild type Art1. While previous models suggested that cargo specificity for ARTs occurs through changes in the permease (Busto et al., 2018; Gournas et al., 2018; Guiney et al., 2016), this is the first demonstration of adaptor-driven specificity. Considering the proximity of loop 3 to the basic patch on Art1 required for recognizing the N-terminal tail of its cargo (figure 1D), it is possible that loop 3 is involved in cargo binding for select permeases. However, a more detailed biochemical and structural analysis, as well as a more complete inventory of Art1 cargos, will be required to define the mechanism through which loop 3 contributes to substrate specificity.

The remaining loop and tail mutants had similar effects on both Mup1 and Can1, suggesting that they are involved in more general regulation of Art1. Although individually deleting the N-terminal tail, loop 1, and loop 2 had no detectable effect when cargo endocytosis was analyzed (figure 2), they did exhibit a reduced ability to associate with the PM (figure 3), and exhibited hypersensitivity to canavanine (figure 2A) and thialysine (figure 2 – supplement 2). Interestingly, combining the deletions in *art1^ΔN,1,2,C^* had a dramatic effect on both Can1 and Mup1 endocytosis (figure 2), as well as Art1 PM association (figure 3), suggesting that each loop and tail makes a minor contribution to Art1 regulation that is exacerbated in combination. Notably, *art1^ΔC^* alone increased Art1 activity. Therefore, the reduced activity of *art1^ΔN,1,2,C^* is likely through the combined effects of the N-terminal tail, loop 1, and loop 2 deletions.

### Phosphoinhibition of Art1

Our data suggest that the previously identified Npr1-targeted residues on Art1 fall within loop 1 (MacGurn et al., 2011). Further, Npr1-independent phosphorylation sites were identified, many of which also occur on loops or the C-tail. Thus, we tested if other kinases regulated Art1 through the phosphorylation of its loops and tails. Consequently, we identified Pho85 as a kinase that affects Art1 function, and further identified Pho80 and Clg1 as the two cyclins that mediate regulation of Art1 by Pho85 (figure 4 and 5). However, regulation by Pho80-Pho85 is indirect, since in *pho80Δ* phosphorylation in loop 1 of Art1 was increased (figure 6A). Additionally, Pho85 is a proline-directed kinase, but changes in phosphorylation at proline-directed sites was not detected in either *clg1Δ* or *pho80Δ.* While there is limited evidence that Pho85 can phosphorylate non-proline-directed sites (Zappacosta et al., 2006), our data cannot rule out that Clg1-Pho85 is acting in a signaling pathway upstream of Art1, rather than directly phosphorylating it, to regulate its activity. In this model, another (yet unidentified) downstream kinase or phosphatase is directly acting upon Art1 (figure 8). Nonetheless, the data here clearly show that Art1 activity is regulated by phosphorylation mediated by Pho80 and Clg1.

**Figure 8.**
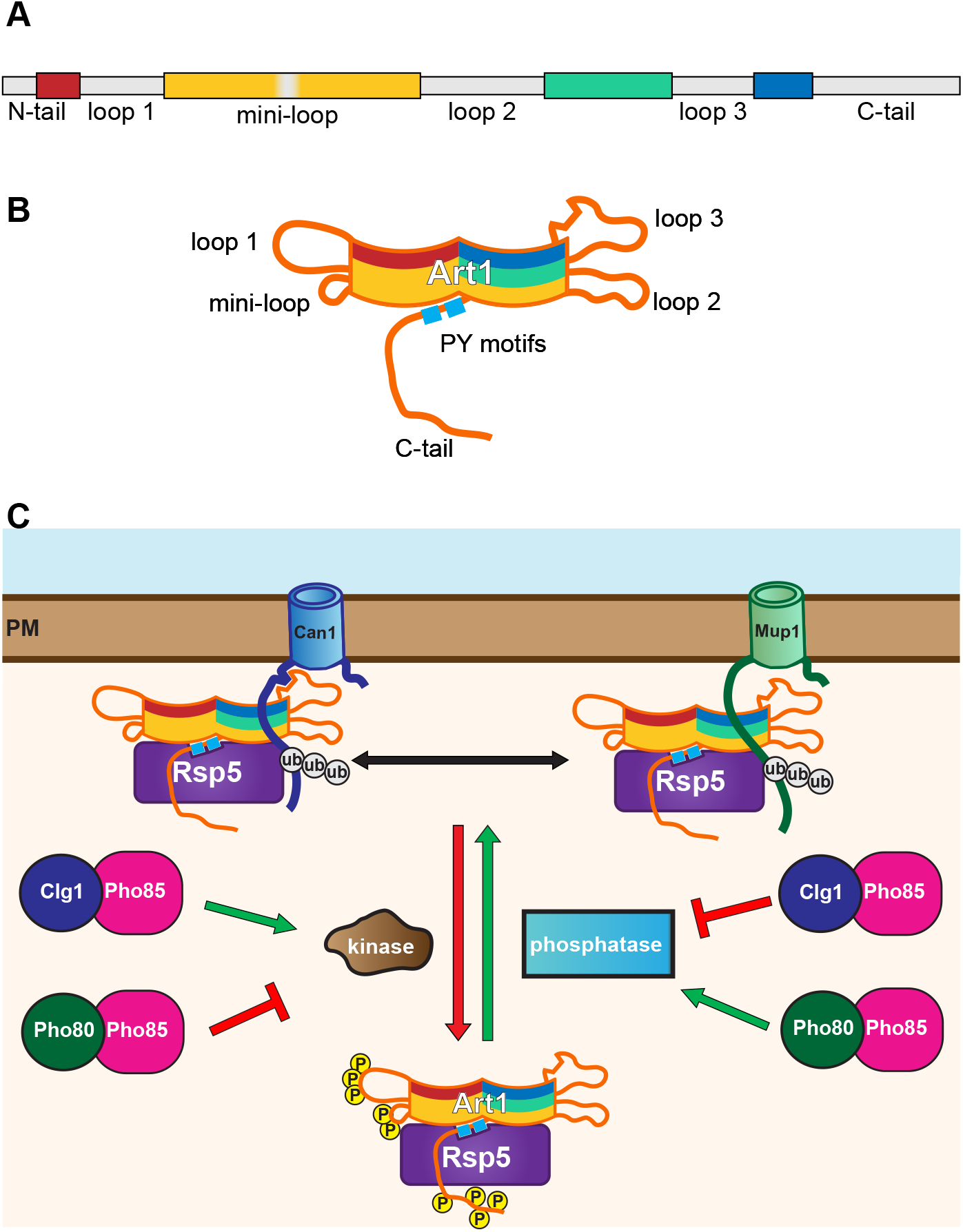
Model of Art1 regulation. (A) Art1 schematic showing the conserved regions predicted to form the arrestin fold (colored), and the loop and tail regions (grey) (B) Cartoon depicting the predicted organization of the conserved arrestin fold and the localization of the loops. Based on the structural model in figure 1F. (C) New modes of Art1 regulation. In addition to the previously identified regulation through Npr1 or Ppz1/2, Art1 phosphorylation/dephosphorylation reactions can be mediated by Clg1-Pho85 and Pho80-Pho85 to regulate its activity. Phosphorylation of Art1 loops and the C-tail regulate its function by affecting its PM localization. When phosphorylated on loop 1, the “mini loop”, and the C-tail, Art1 remains on the cytosol or associated with the Golgi, rendering it inactive. Upon dephosphorylation Art1 can associate with the PM, bringing it into proximity with its substrate cargos resulting in ubiquitylation and endocytosis. Additionally, loop 3 is required for recognition of a subset of cargos.

The function of Clg1 is poorly defined. Despite belonging to the Pcl1/2 subfamily (Matsumoto and Wickner, 1993), which usually is involved in cell cycle control, mutating *CLG1* alone results in no detectable defect in cell cycle progression, consistent with its constitutive expression throughout the cell cycle (Measday et al., 1997). More recently, there is evidence that Clg1 phosphorylates and degrades Sic1 and therefore promotes autophagy (Yang et al., 2010). Notably, amino acid homeostasis is altered in *clg1Δ* (Mülleder et al., 2016). Thus, similar to the TORC1-Npr1 pathway which inactivates Art1 during starvation conditions (MacGurn et al., 2011), Clg1 may regulate Art1 in response to changes in nutrient levels. In this model, low nutrient/amino acid levels would cause Clg1 activation, which would in turn inactivate Art1, increasing the abundance of permeases on the PM. Identification of the input(s) that regulate Clg1 as well as a complete inventory of its downstream targets will be critical to understanding this signaling pathway.

In comparison, Pho80 function is much better defined. Pho80 is a critical component of the phosphate sensing pathway in yeast (Kaffman et al., 1994), is required for survival under various stress conditions (Huang et al., 2002), and is involved in Rim15-mediated nutrient sensing (Wanke et al., 2005). Although Art1 has not yet been implicated directly in either the phosphate response or Rim15 pathway, both are important metabolic pathways and modulation of PM permeases to control nutrient levels may be required for either cellular homeostasis or survival.

The increased phosphorylation of loop 1 observed in *pho80Δ* occurs at residues distinct from those identified as phosphorylated by Npr1 (figure 6A, MacGurn et al., 2011). However, the phenotypes are similar. Phosphorylation mimetic mutations at S92/T93 result in Art1 inactivation and a failure to associate with the PM (figure 6 and 7). Although the cellular inputs are different, the mechanism through which they attenuate Art1 function may be the same. In the structural model of Art1, loop 1 would extend distally from the N-terminal arrestin domain (figure 1E, figure 1 – supplement 2). Thus, phosphorylation at any number of residues on this loop seem to prevent PM localization. This suggests that this region of Art1 interacts with either a PM resident protein or the lipids that constitute the PM, and that phosphorylation disrupts this interaction. Interestingly, the non-conserved “mini-loop,” which contains residues 238-241, is predicted to occur at a turn between two β-strands that is spatially adjacent to loop 1. Therefore, this region may also be involved in Art1 binding to the PM or PM proteins, or at least phosphorylation of this region can disrupt this binding. While the structural model did not include the C-terminal tail, considering the effect on cargo endocytosis in the Asp mutants, and the effect on Art1 PM localization in the absence of the tail, the C-terminal tail likely regulates Art1 similarly, although it is currently unclear if this occurs through a similar or distinct membrane binding region. Nonetheless, one aspect of Art1 regulation is its ability to properly localize to the membrane containing its substrates, which occurs via phosphorylation and dephosphorylation. Notably, it is currently unclear if Art1 is phosphorylated and inactivated in the cytosol to prevent membrane association, or if PM-associated Art1 is phosphorylated to disengage it from the membrane and decrease its activity.

Recently, the Ppz phosphatases have been identified as regulators of Art1 (Lee et al., 2019). Intriguingly, one of the targets of Ppz1/2 is T93. Therefore, the possibility exists that either Pho80 or Clg1 regulate Art1 by acting on the Ppz proteins. Interestingly, T245 and T795 were also identified as Ppz targets that affect Art1 function. While these residues were not identified in our study, T245 falls within the “mini-loop” of Art1, and T795 is on the C-tail. Thus, phosphorylation in these regions, regardless of the specific input, all regulate Art1.

This work paints a more complete picture of the molecular complexity underlying Art1 regulation during selective cargo ubiquitylation and endocytosis (figure 8). In order for cargo to be sorted to the vacuole, it must first bind substrate, changing its conformation, and allowing Art1-Rsp5 mediated binding and ubiquitination (Busto et al., 2018; Gournas et al., 2017, 2018; Guiney et al., 2016). Art1 binding to specific cargos can be modulated by loop 3 in Art1. Further, ubiquitylation of cargo can be enhanced by Art1 PM association, which is controlled via phosphorylation. Multiple different signaling pathways, which include Clg1-Pho85, Pho80-Pho85, and TORC1/Npr1, affect Art1 phosphorylation in response to a diverse array of environmental cues. Therefore Art1, and likely the other ART proteins, is able to be finely tuned to remodel the composition of the PM and in turn, cellular homeostasis.

## MATERIALS AND METHODS

### Yeast strains and growth conditions

All yeast strains used in this study were generated from SEY6210 (Robinson et al., 1988) and are listed in extended data file 3. Genetic knockouts were generated by replacing the entire open reading frame of the gene using PCR-mediated gene replacement (Goldstein and McCusker, 1999; Longtine et al., 1998; Wach et al., 1994), and multiple mutants were generated by crossing, sporulation, and tetrad dissection (Sherman, 2002). Chromosomally tagged strains were generated by PCR-mediated tagging using pFA6a-mCherry-kanMX (Malcova et al., 2016) to tag *VPH1,* pFA6a-GFP (Longtine et al., 1998) to tag *MUP1,* and pFA6a-LAP-eGFP-HIS3 (Adell et al., 2017) to tag *CAN1.* pRS305- and pRS306-based vectors (Sikorski and Hieter, 1989) were integrated after linearization with AflII or StuI, respectively, unless the inserted gene contained additional AflII or StuI sites, in which case the plasmid was linearized by PCR amplification.

Cells were grown at 26°C in synthetic media [0.17% (w/v) yeast nitrogen base, 0.5% (w/v) ammonium sulfate, 2% (w/v) glucose, and supplemented with adenine, uracil, Arg, Lys, Thr, Tyr, His, Leu, and Trp; unless otherwise specified. For acute YPD treatment, cells were grown in synthetic media, washed with dH_2_O, then resuspended in YPD [1% (w/v) yeast extract, 1% (w/v) peptone, 2% (w/v) glucose] and incubated for the indicated time. For growth on canavanine, solid synthetic media (containing 2% agar) lacking Arg was supplemented with the indicated concentration of canavanine (Sigma) after autoclaving. Similarly, for growth on thialysine, solid synthetic media lacking Lys was supplemented with the indicated concentration of thialysine (Sigma) after autoclaving. For grow assays, 1:10 serial dilutions were spotted onto the indicated media using a frogger pinning apparatus and incubated at 26°C unless otherwise specified.

### Plasmids

All plasmids used in this study are listed in extended data file 4 and were generated using standard procedures and conditions. All restriction endonucleases were purchased from New England Biolabs. Plasmids containing *ART1* included 536 bp of the 5’ UTR as the promoter and 502 bp of the 3’ UTR as the terminator. The HTF tag (MacGurn et al., 2011), consisting of a 6x <u>H</u>is tag, a <u>T</u>EV cleavage site, and a 3x <u>F</u>LAG tag, contains a RIPGLINRGS linker between the C-terminus of Art1 and the 6x His tag. The mNeonGreen (mNG) tag (Shaner et al., 2013) adds a RIPGLINIF linker between Art1 and NG. Both tags were added to *ART1* containing plasmids using overlap extension (Ho et al., 1989). All *ART1* point mutants were generated using overlap extension. pEG166 included 596 bp of the MUP1 5’ UTR as the promoter and 279 bp of the 3’ UTR as the terminator, and introduced the HTF tag as an EagI-SacII fragment, containing a RPRIPGLINRGS linker between the C-terminus of Mup1 and the 6x His tag. pMB0268 was generated by inserting the *CLG1* coding sequence immediately 3’ of the *GPD (TDH3)* promoter. pMB0408 was generated by removing the *CEN* sequence from pRS317 via PCR, then ligating P_*GPD*_CLG1 into the MCS. pMB0239 was generated by ligating the *SEC7* coding sequence with 527 bp of the 5’ UTR and 303 bp of the 3’ UTR into pRS305. mCherry was inserted on the 5’ end of *SEC7* by overlap extension and contains an SSGSGSS linker between mCherry and Sec7. All plasmids were verified by sequencing.

### Endocytosis assays

For Mup1-GFP endocytosis, cells were grown at 30°C in synthetic media lacking Met, and were diluted multiple times to ensure that the culture never exceeded ODβ00 = 1 for over 24 hrs. The indicated concentration of Met was added to initiated endocytosis, and cells were collected after 60 min for further analysis. Can1-GFP endocytosis was performed similarly, except the synthetic media lacked Arg, and endocytosis was initiated with the indicated concentration of Arg.

### Protein extraction and immunoblotting

5 OD_600_ equivalents of yeast were incubated in 10% (v/v) TCA on ice overnight. Cells were pelleted, washed with cold acetone, and dried. Cells were then resuspended in 50 μL urea cracking buffer [50 mM Tris-HCl pH 7.5, 8 M urea, 2% (w/v) SDS, 1 mM EDTA], lysed with glass beads by vortexing for 5 min at room temperature, then incubated at 42°C for 5 min. 50 μL 2x sample buffer [150 mM Tris-HCl pH 6.8, 7 M urea, 10% (w/v) SDS, 24% glycerol, 10% (v/v) β-mercaptoethanol, bromophenol blue] was added and the samples were vortexed again for 5 min at room temperature, then incubated at 42°C for 5 min. Proteins were separated by SDS-PAGE and transferred onto 0.45 μm nitrocellulose membranes (GE Healthcare) in transfer buffer (25 mM Tris, 192 mM glycine, 10% methanol). When analyzing Mup1-GFP or Can1-GFP, transfer buffer was supplemented with 0.06% SDS.

Membranes were blocked in blocking buffer [5% (w/v) fat-free milk in TBST (20 mM Tris-HCl pH 7.5, 150 mM NaCl, 0.05% Tween-20)] for 60 min. Primary antibodies were diluted in blocking buffer and incubated for 4 hours at room temperature or overnight at 4°C. Membranes were washed 3x for 10 min in TBST. Secondary antibodies were diluted in blocking buffer and incubated for 60 min at room temperature, then membranes were washed 3x for 10 min in TBST. Membranes were scanned using an Odyssey CLx imaging system (Li-Cor Biosciences) and analyzed using ImageStudio Lite version 5.2 (Li-Cor Biosciences) or Image J (National Institutes of Health).

The primary antibodies used were mouse anti-FLAG M2 (Sigma-Aldrich F1804) 1:5000, rabbit anti-glucose-6-phosphate dehydrogenase (G6PDH, Sigma-Aldrich A9521) 1:50,000, mouse anti-GFP (Roche 11814460001) 1:5000 for detecting Can1-GFP, mouse anti-GFP (Santa Cruz Biotechnology sc-9996) 1:500 for detecting Mup1-GFP, and rabbit anti-Rsp5 (Emr Lab) 1:10,000. The secondary antibodies used were goat anti-mouse 800 CW (Li-Cor Biosciences 926-32210) 1:10,000 and goat anti-rabbit 680 LT (Li-Cor Biosciences 926-68021).

### Fluorescence microscopy

Microscopy was performed using a DeltaVision RT system (Applied Precision, Issaquah, WA), equipped with a DV Elite CMOS camera, a 100 x objective, and a DV Light SSI 7 Color illumination system with Live Cell Speed Option (FITC for GFP and TRITC for mCherry/RFP). Image acquisition and deconvolution were performed using the provided DeltaVision software (softWoRx 6.5.2; Applied Precision, Issaquah, WA).

Quantification of Art1-mNG was conducted using a custom Python pipeline. Source code is available at github.com/elguiney/YeastMorpholgyPypeline, commit 4fd7685. Briefly, edge detection using a Laplace of Gaussian kernel was used to define cell boundaries from bright field images. For each cell, Golgi masks were defined based on a maximum intensity z-projection of the deconvolved mCherry-Sec7 signal, thresholded with Otsu’s method. Because Art1-mNG fluorescence at the Golgi was often directly adjacent to the mCherry-Sec7 marker, the Golgi mask regions were expanded (by ensuring that the local radius of the mask was always >= 7 pixels). A cell cortex mask was defined as the region within 10 pixels of the cell border on the inside, and 5 pixels on the outside, that did not overlap with either a Golgi mask or an adjacent cell. Analysis was run in batch on all images from three independent biological replicates. After cell identification and mask generation, cells were cropped and randomized, and displayed for blinded visual inspection; cells where the cell boundary or the Golgi or cell cortex masks were incorrectly identified were rejected from further analysis (typically fewer than 10% of cells were rejected). Finally, for each cell Art1-mNG enrichment in the Golgi and cell-cortex mask regions was calculated by subtracting the cytoplasmic background Art1-mNG fluorescence (as defined by regions in a cell not covered by a Golgi or cell-cortex mask); and then normalized to total Art1-mNG fluorescence in the cell.

### Immunoprecipitation

Anti-FLAG immunoprecipitation of Art1-HTF was performed by collecting 25 OD_600_ equivalents of yeast in mid-log (OD_600_ = 0.5 – 0.8) growth, followed by a wash in 20 mL of 10 mM Tris-HCl pH 8.0, 5 mM EDTA, 10 mM NEM, 1 mM PMSF, 10 mM β-glycerophosphate, 5 mM NaF. Cell pellets were collected and stored at −80°C for at least 12 hours. Cell pellets were then thawed on ice and resuspended in 500 μL lysis buffer (50 mM Tris-HCl pH 7.4, 150 mM NaCl, 5 mM EDTA, 1 mM PMSF, 10 mM NEM, 1x cOmplete protease inhibitor cocktail (Roche), 10 mM β-glycerophosphate, 5 mM NaF). Cells were lysed by vortexing with 0.5 mm YZB zirconia beads (Yasui Kikai) for 30 sec three times, with 30 sec incubations on ice in between. 500 μL lysis buffer containing 0.1 % Igepal CA-630 (USB corporation) was added, and lysate incubated on ice for 10 min. Unbroken cells and the beads were removed by centrifugation at 500 x *g* for 10 min at 4°C, then lysates were cleared by centrifugation at 16,000 x *g* for 15 min at 4°C. The cleared lysates were incubated with preequilibrated EZview Red anti-FLAG M2 Affinity Gel (Sigma) at 4°C while rotating for 2 hrs. The resin was washed in lysis buffer 3 times, and the bound proteins eluted by incubating in elution buffer (100 mM Tris-HCl pH 8.0, 1% SDS, 10 mM DTT) at 95°C for 10 min.

### Mass spectrometry

For analysis of Art1 phosphorylation, cells were grown to mid-log in synthetic media supplemented with adenine, uracil, Asp, Ile, Leu, Phe, Thr, Trp, Tyr, Val, His, and Pro, and either light Arg and Lys or heavy Arg (^13^C_6_ ^15^N_4_, Sigma) and Lys (^13^C_6_ ^15^N_2_, Sigma). Art1-HTF was immunoprecipitated as described above, except 250 OD_600_ equivalents were harvested and the lysis buffer additionally contained 1x PhosSTOP phosphatase inhibitor (Roche). Eluates were reduced with 10 mM DTT for 15 min at room temperature, then alkylated with 20 mM iodoacetamide. Heavy and light samples were mixed 1:1 and precipitated by adding 3 volumes of 50% acetone, 49.9% ethanol, 0.1% acetic acid. Proteins were resuspended in 8 M urea, 50 mM Tris-HCL pH 8.0, diluted with 3 volumes of dH_2_O, and digested overnight with 1 g trypsin at 37°C.

Phosphopeptides were enriched using Fe-NTA as described elsewhere (Bastos de Oliveira et al., 2018). Data were analyzed using both the Sorcerer-SEQUEST data analysis pipline (Bastos de Oliveira et al., 2018) and MaxQuant for phosphopeptide identification, quantitation and phosphorylation site localization. MS/MS spectra were also manually inspected for MS2 quality and to confirm phosphorylation site localization. MaxQuant site localization score is included in the supplementary data. Scatter plots were generated using a data analysis package courtesy of Haiyuan Yu (Cornell University).

### Bioinformatic analysis

For analysis of Art1 conservation, homologs were retrieved from the NCBI refseq database with BLAST, sequences annotated as ‘partial read’ removed. Art1 homologs were defined as proteins with a BLAST E-value less than 0.1, and whose top BLAST hit in the *Saccharomyces cerevisiae* proteome was also Art1. Likewise, Art2/8, Art3/6 and Art4/5/7 homologs were defined as proteins with a BLAST E-value less than 0.1 when Art2, Art3, or Art4 were used as query, respectively, and whose top BLAST hit in the *Saccharomyces cerevisiae* proteome was also Art2 or Art8, Art3 or Art6, or Art4, Art5 or Art7 respectively. Homologs were aligned with mafft, using the -linsi option (--maxiterate 1000 --localpair); phylogenetic trees were computed with FastTree (Price et al., 2010), using the Le-Gascuel subsitition model and pseudocounts (options -lg and -pseudo); Art1, Art3/6, and Art4/5/7 multiple sequence alignments were rooted to TXNIP from *Homo sapiens,* and Art2/8 multiple sequence alignment was rooted to Uniprot ID Q6CG62 from *Yarrowia lipolytica*. Conservation was computed with the rate4site algorithm (Mayrose et al., 2004; Pupko et al., 2002), using the Le-Gascuel substitution model and Bayesian rate inference (options -Ml and -ib); conservation scores shown in Figure 1, and Figure 1 supplements 4–7 are the lower bound of the 25-75% confidence interval for the estimated evolutionary rate parameter, filtered with a 5 residue sliding window to provide a conservative estimate of relative conservation across gappy alignments. Structural modeling of S. *pombe* Any1p and Art1^ΔN12C^ was performed using the iTASSER webserver (Roy et al., 2010; Yang et al., 2010; Zhang, 2008).

## ACKNOWLEDGEMENTS

The authors would like to thank Richa Sardana, Sho Suzuki, and Jeff Jorgensen for critical reading of the manuscript, and Lu Zhu for constructing the pRS307 plasmid. M.G.B. was supported by a Sam and Nancy Fleming Research fellowship. This work was supported by a Cornell University Research Grant to S.D.E and NIH grant R00GM101077 to J.A.M.

---

**Figure 1 – supplement 1.**
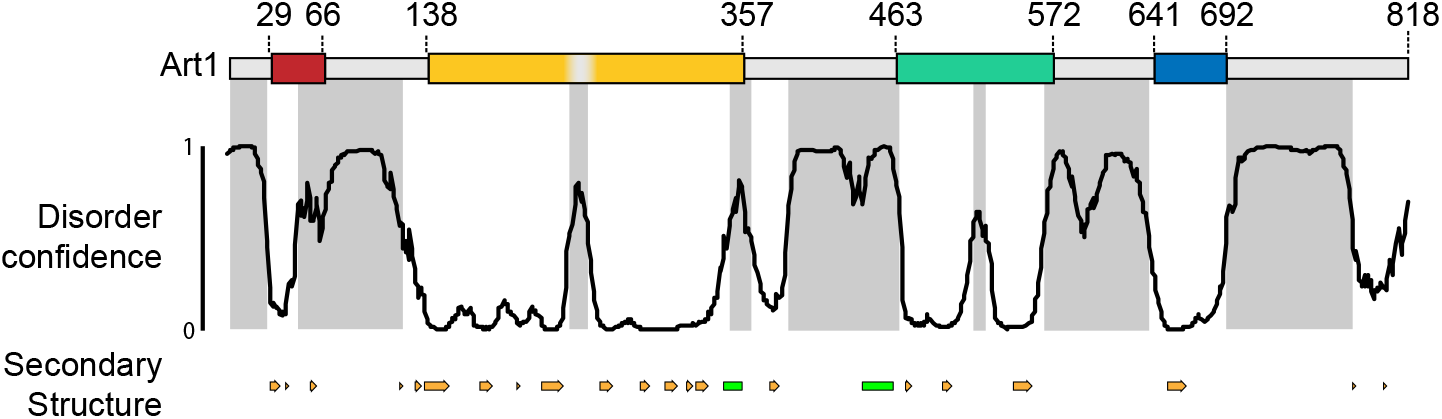
Art1 loops are predicted to be disordered. Disorder and secondary structure predictions for Art1. (top) Art1 schematic, as in figure 1D. (middle) Solid black line, Disorder confidence predicted by DISOPRED3 server. Grey regions, predicted disordered regions. (bottom) Secondary structure predicted by PSIPRED server, orange arrows = β-sheets, green boxes = α-helices.

**Figure 1 – supplement 2.**
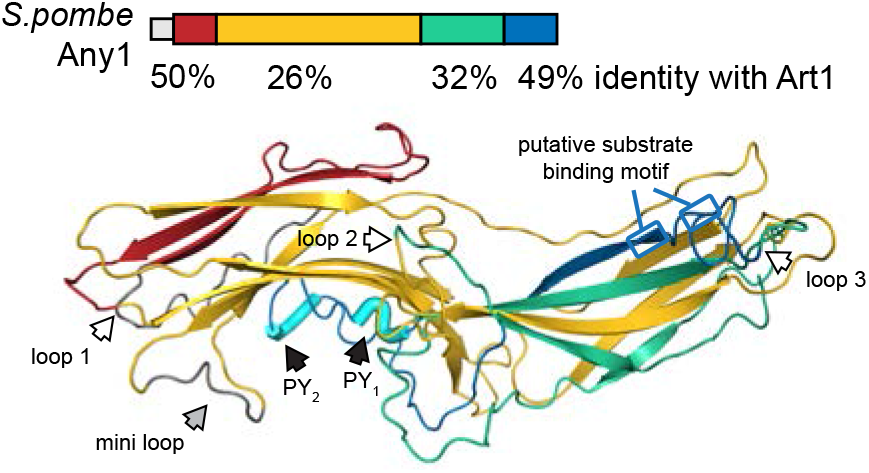
art1^ΔN,1,2,C^ is predicted to form an arrestin fold. (top) Schematic of *S.pombe* Any1/Arn1, with the indicated % identities compared to the corresponding conserved regions of Art1. (bottom) Structural model of Any1. The position of the predicted loop locations in Art1, the PPXY motifs, and the substrate binding residues are indicated.

**Figure 1 – supplement 3.**
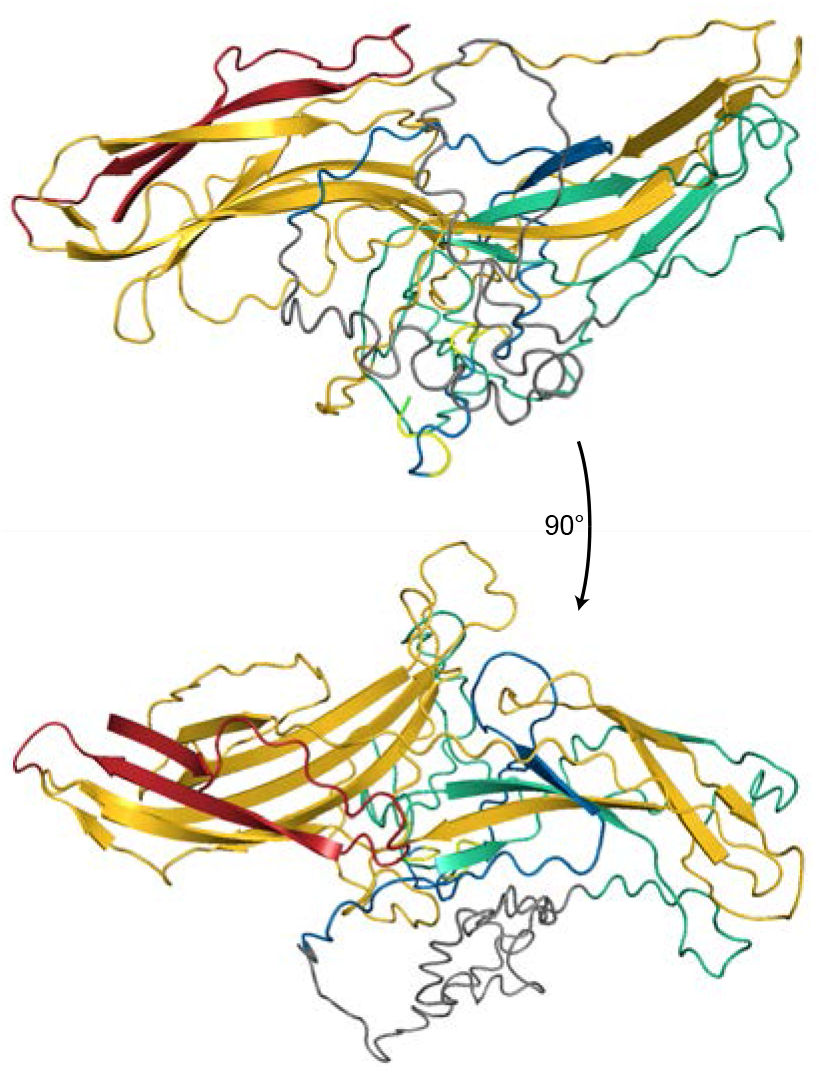
art1^ΔN,2,C^ is predicted to form an arrestin fold. Structural model of art1^ΔN,1,2,C^. Core regions are colored as in figure 1. Loop 3 is shown in grey

**Figure 1 – supplement 4.**
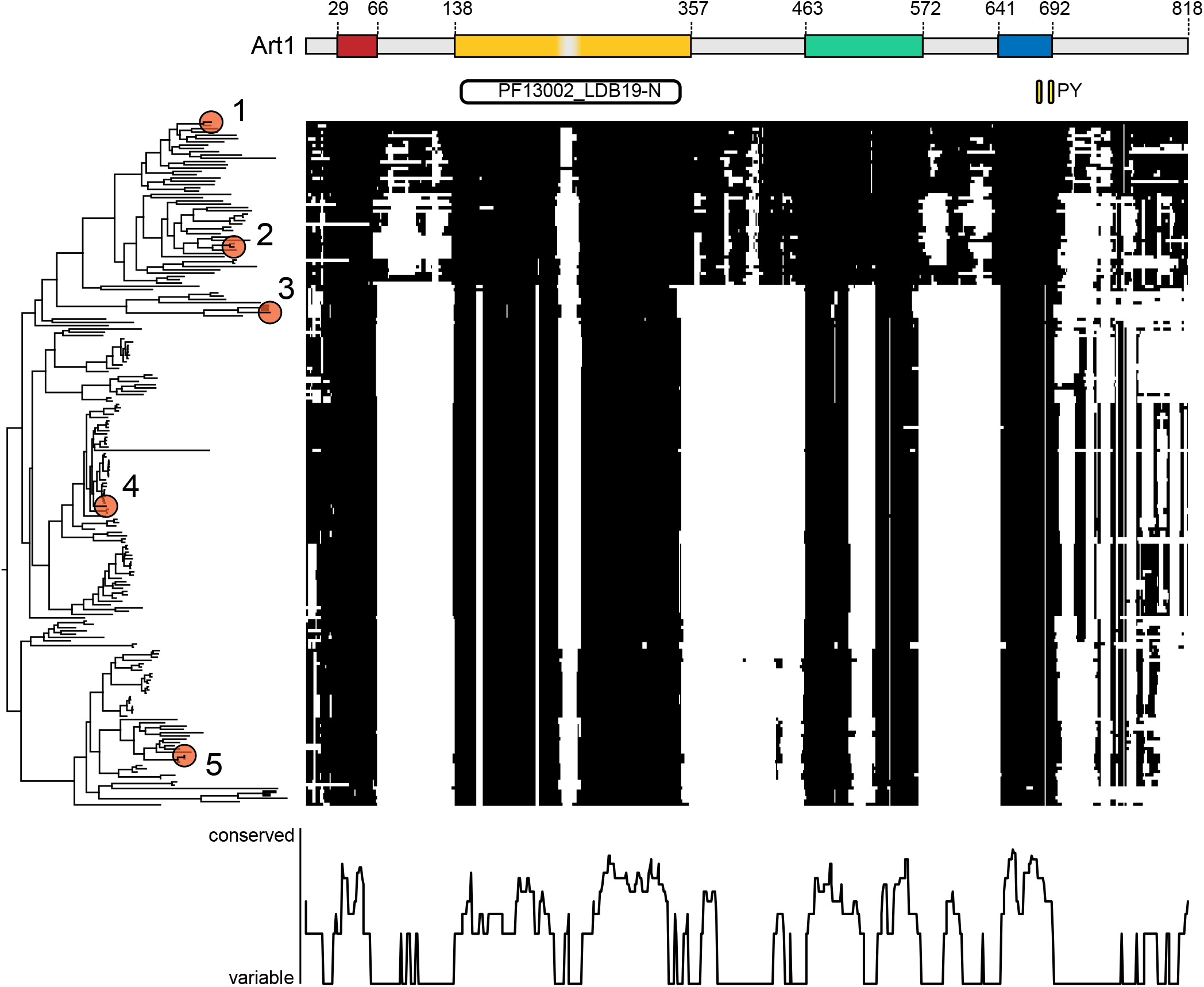
Art1 conservation in fungi. Summary plot of Art1 homolog multiple sequence alignment. Black regions indicate residues that align to Art1, white regions indicate gaps. A gene tree is displayed on the y axis with the following homologs indicated by numbered circles: (1) *Saccharomycse cerevisiae* Art1 (uniprot ID Q12502) (2) *Candida albicans* (Q59NC8) (3) *Schizosaccharomyces pombe* Any1p (O60150) (4) *Aspergilus nidulans* (Q5BHC4) (5) *Neurospora crassa* (Q7S1Q1). (bottom) Conservation of Art1 residues plotted on the x axis, as in figure 1D.

**Figure 1 – supplement 5.**
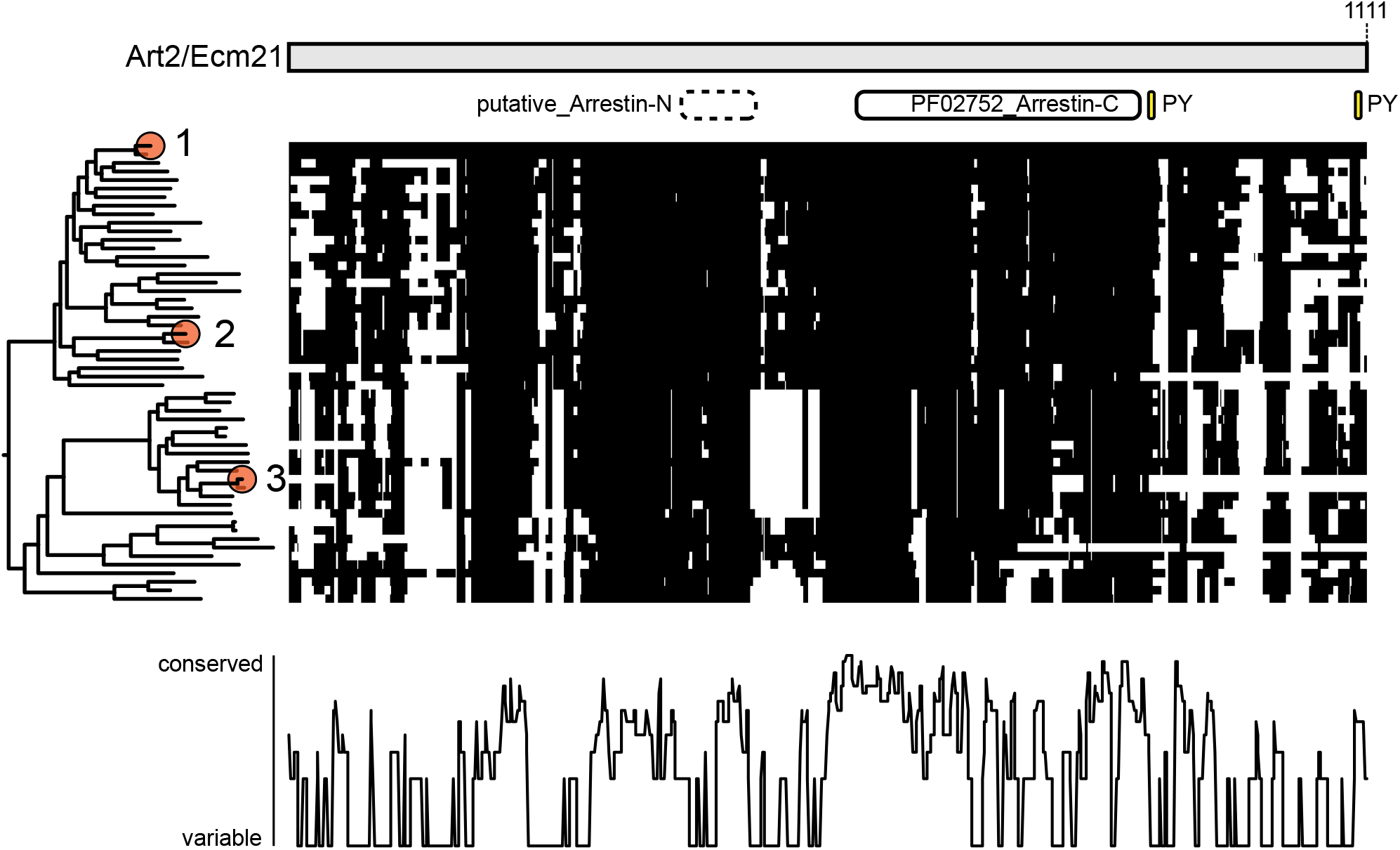
Art2/8 conservation in fungi. Summary plot of Art2 homolog multiple sequence alignment. Black regions indicate residues that align to Art2, white regions indicate gaps. A gene tree is displayed on the y axis with the following homologs indicated by numbered circles: (1) *Saccharomyces cerevisiae* Art2/Ecm21 (uniprot ID P38167), (2) *Saccharomyces cerevisiae* Art8/Csr2 (Q127374) (3) *Candida albicans* (A0A1D8PET4). (bottom) Conservation of Art2 residues plotted on the x axis.

**Figure 1 – supplement 6.**
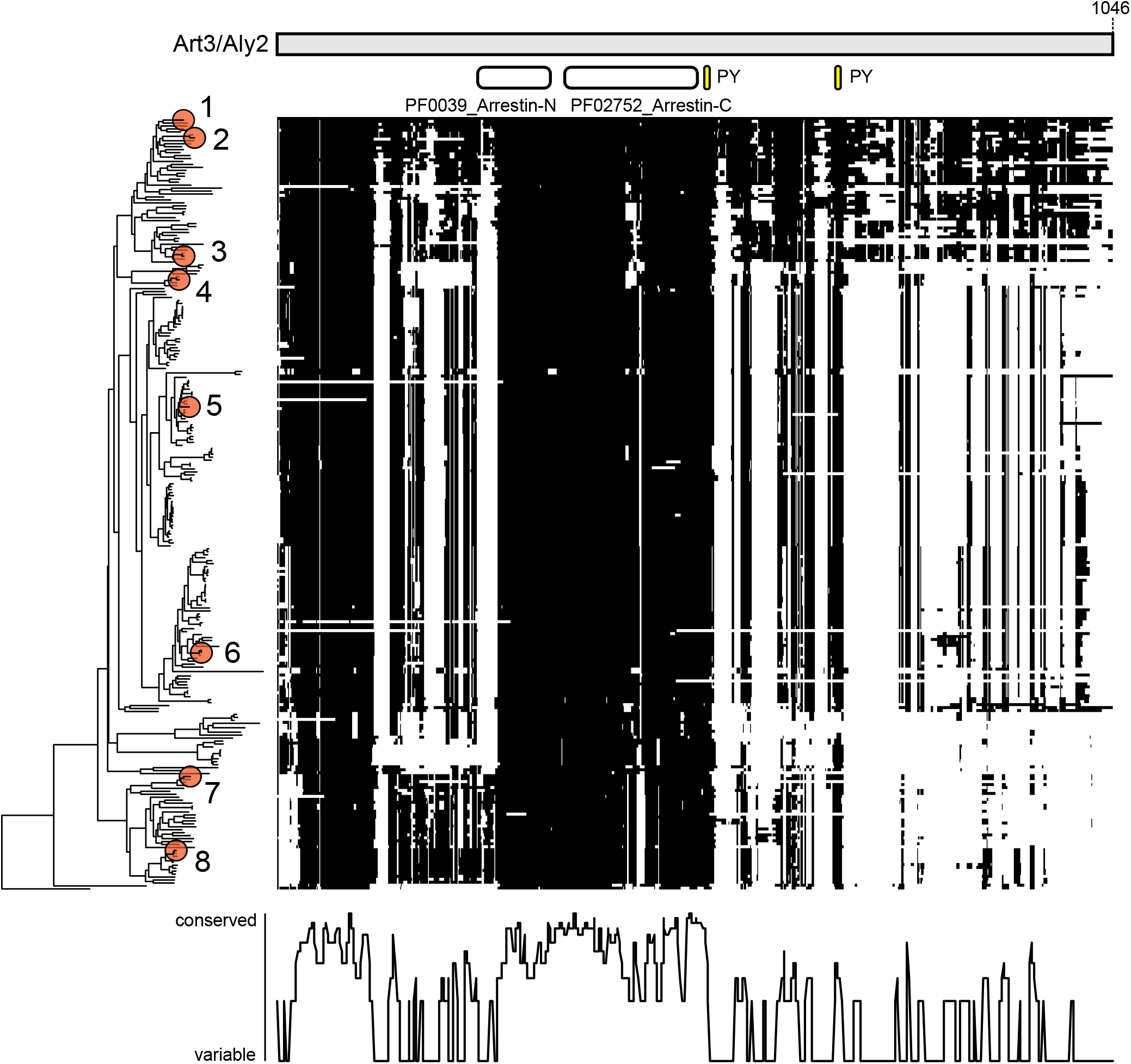
Art3/6 conservation in fungi. Summary plot of Art3 homolog multiple sequence alignment. Black regions indicate residues that align to Art3, white regions indicate gaps. A gene tree is displayed on the y axis with the following homologs indicated by numbered circles: (1) *Saccharomyces cerevisiae* Art3/Aly2 (uniprot ID P47029), (2) *Saccharomyces cerevisiae* Art6/Aly1 (P36117), (3) *Candida Albicans* (A0A1D8PQ35), (4) *Schizosaccharomyces pombe* (O74798), (5) *Aspergillus nidulans* (Q5BEE1), (6) *Neurospora crassa* (Q7SB91), (7) *Ustilago maydis* (A0A0D1E8U6), (8) *Cryptococcus neoformans* (Q55SH7). (bottom) Conservation of Art3 residues plotted on the x axis.

**Figure 1 – supplement 7.**
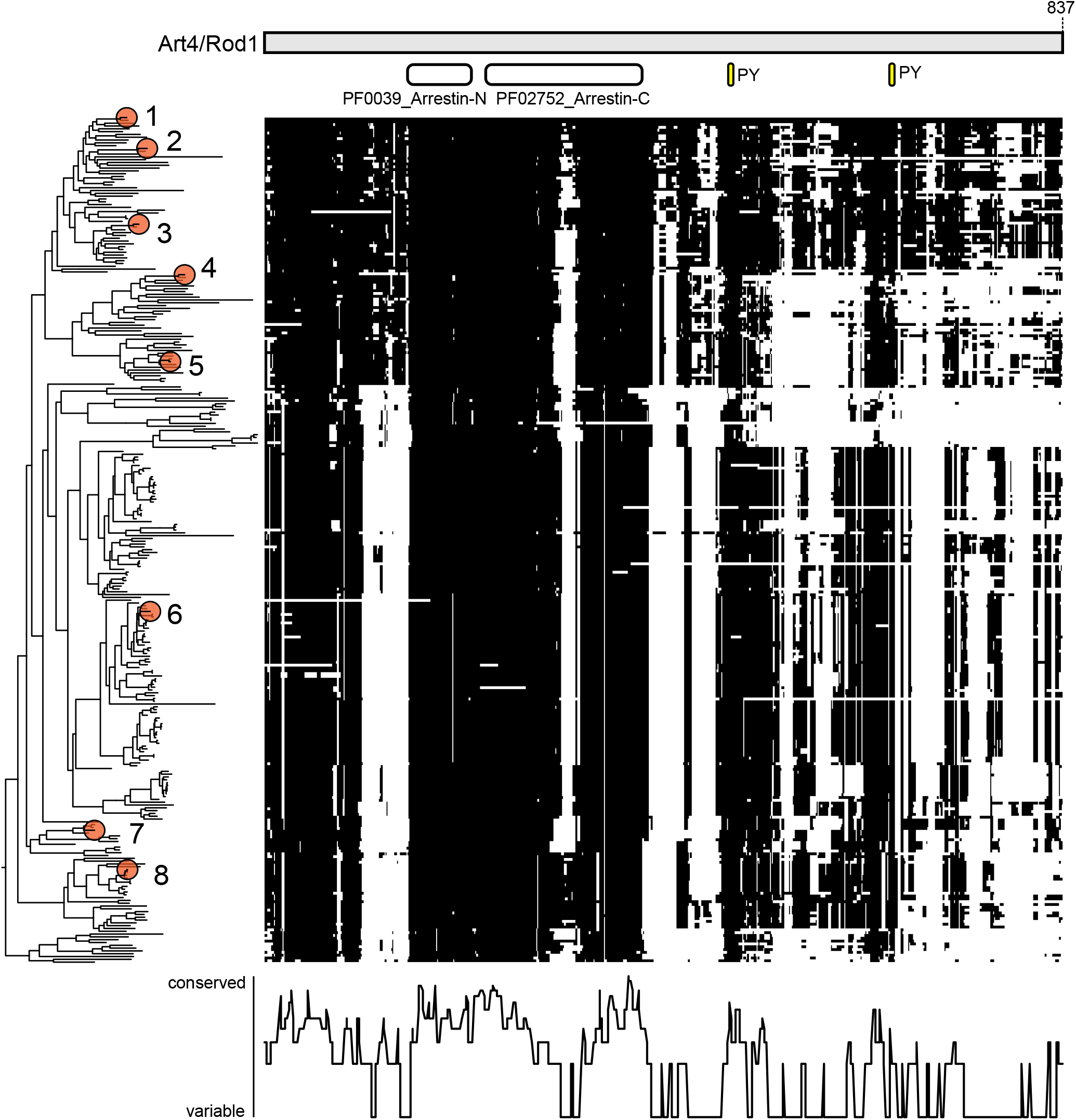
Art4/5/7 conservation in fungi. Summary plot of Art4 homolog multiple sequence alignment. Black regions indicate residues that align to Art4, white regions indicate gaps. A gene tree is displayed on the y axis with the following homologs indicated by numbered circles: (1) *Saccharomyces cerevisiae* Art4/Rod1 (uniprot ID Q02805), (2) *Saccharomyces cerevisiae* Art7/Rog3 (P43602), (3) *Candida albicans* Rod1 (A0A1D8PGG1), (4) *Saccharomyces cerevisiae* Art5 (P53244), (5) *Candida albicans* (A0A1D8PCZ8), (6) *Aspergillus nidulans* creD (Q6SIF1), (7) *Schizosaccharomyces pombe* (Q09729), (8) *Cryptococcus neoformans* (Q55W94). (bottom) Conservation of Art4 residues plotted on the x axis.

**Figure 2 – supplement 1.**
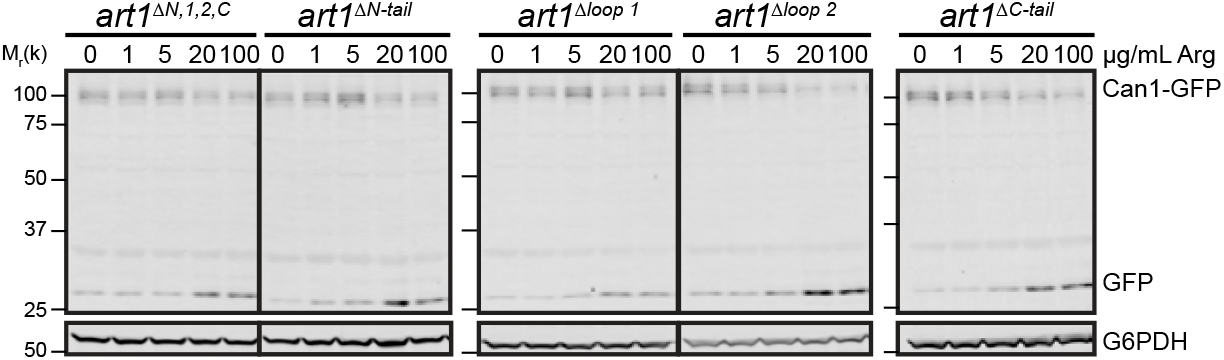
Cargo endocytosis in Art1 loop and tail mutants. Immunoblot analysis of Can1-GFP endocytosis induced with the indicated concentration of Arg for 60 min, in the indicated strains.

**Figure 2 – supplement 1.**
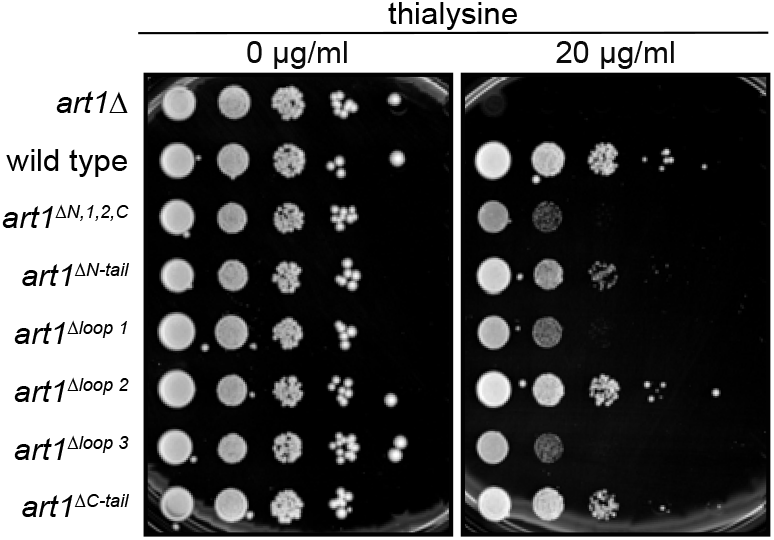
Art1 loop and tail mutant growth on thialysine. Serial dilutions of the Art1 tail and loop mutant strains spotted on synthetic media containing the indicated concentration of thialysine and incubated at 26°C.

**Figure 2 – supplement 1.**
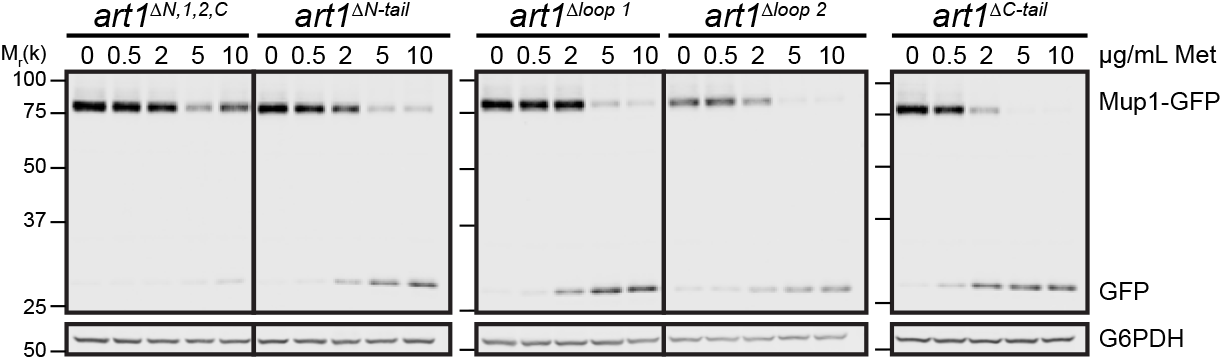
Cargo endocytosis in Art1 loop and tail mutants. Immunoblot analysis of Mup1-GFP endocytosis induced with the indicated concentration of Met for 60 min, in the indicated strains.

**Figure 3 – supplement 1.**
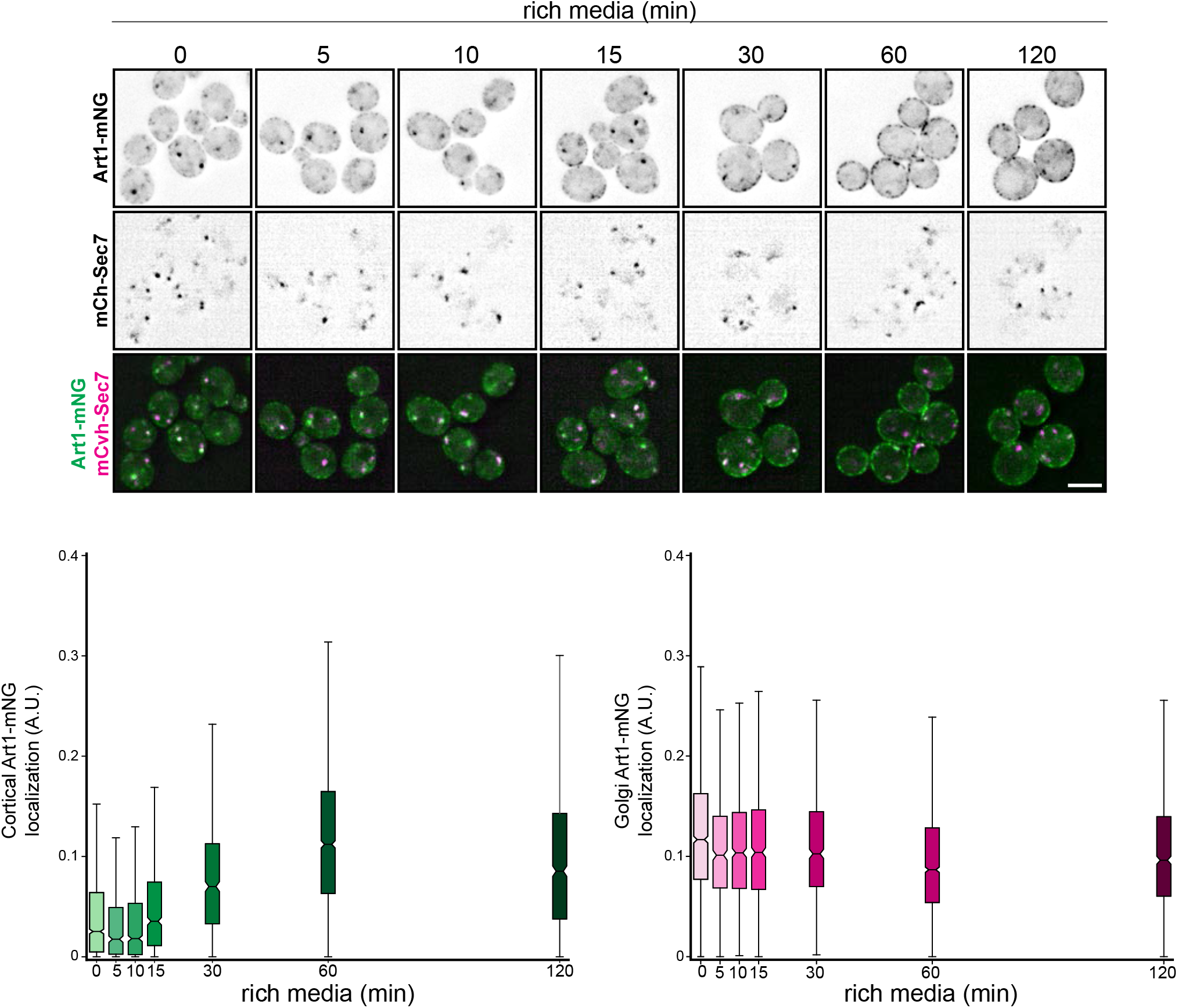
Art1-mNG translocates to the PM after shift to rich media. Art1-mNG localization in minimal media and after shifting to rich media for the indicated times. Quantification of cortical (bottom left) and Golgi (bottom right) Art1-mNG localization at the indicated timepoints. Box and whisker plots show median (horizontal line) and bootstrapped 95% confidence interval of median (notch), 25 and 75% (box) and 1.5*interquartile range (whiskers). From 3 independent experiments; n > 500 cells measured at each timepoint.

**Figure 4 – supplement 1.**
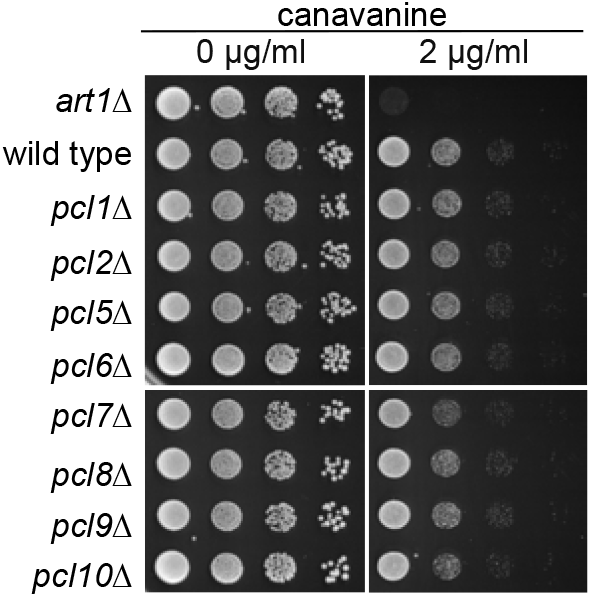
Cyclin mutant growth on canavanine. Serial dilutions of the indicated null mutants spotted on synthetic media containing the indicated concentration of canavanine and incubated at 26°C.

**Figure 4 – supplement 2.**
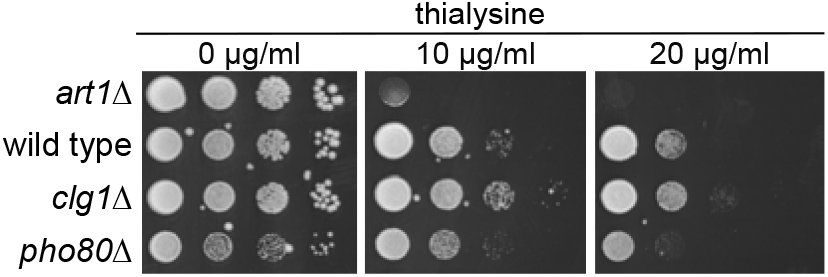
Cyclin mutant growth on thialysine. Serial dilutions of the indicated null mutants spotted on synthetic media containing the indicated concentration of thialysine and incubated at 26°C.

**Figure 4 – supplement 3.**
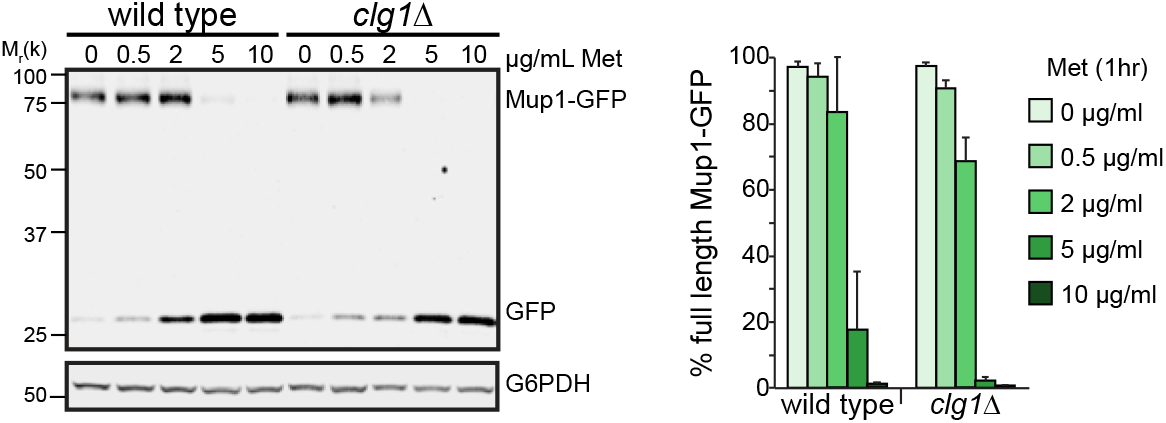
Mup1-GFP degradation in *clg1Δ.* Immunoblot analysis of Mup1-GFP endocytosis induced with the indicated concentration of Met for 60 min in wild type or *clg1Δ* yeast. Band intensities from three biological replicates were quantified and expressed as the mean *%* full length Mup1-GFP remaining. Bars indicate 95% confidence intervals.

**Figure 4 – supplement 4d.**
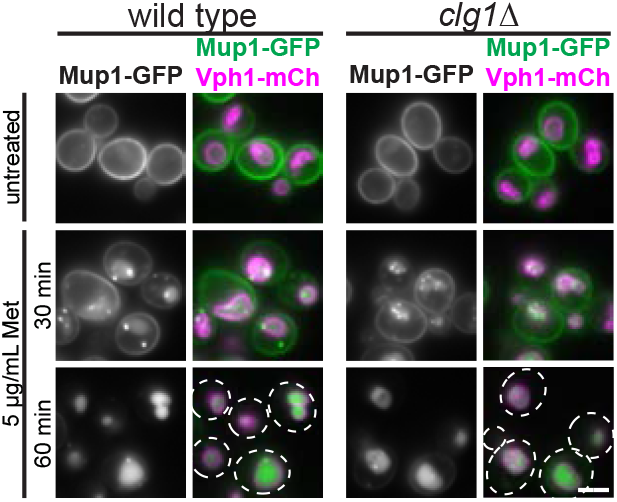
Mup1-GFP endocytosis in *clg1Δ.* Fluorescence microscopy of wild type or *clg1Δ* cells expressing Mup1-GFP, with and without inducing endocytosis by treating with 5 μg/mL Met for 30 and 60 min. Bar = 2 μm

**Figure 4 – supplement 5.**
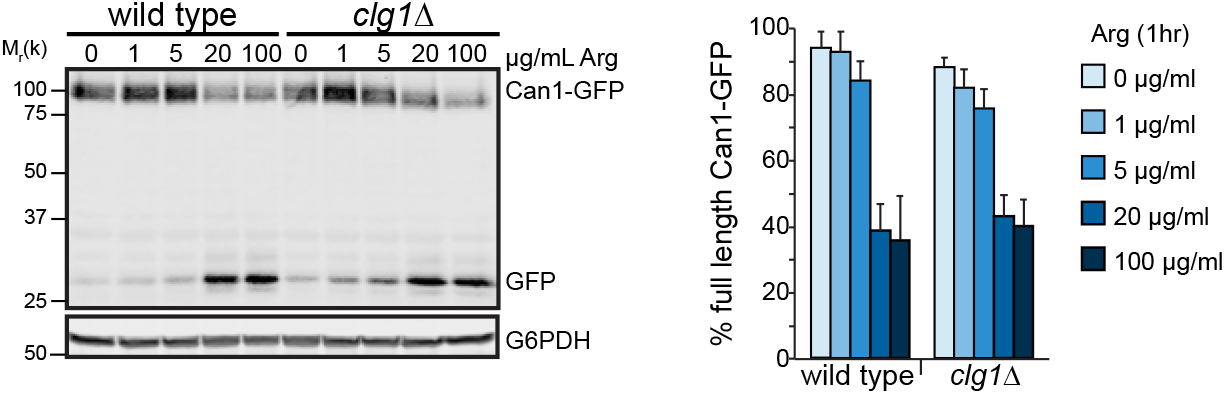
Can1-GFP degradation in *clg1Δ.* Immunoblot analysis of Can1-GFP endocytosis induced with the indicated concentration of Can for 60 min in wild type or *clg1Δ* yeast. Band intensities from three biological replicates were quantified and expressed as the mean % full length Can1-GFP remaining. Bars indicate 95% confidence intervals.

**Figure 4 – supplement 6.**
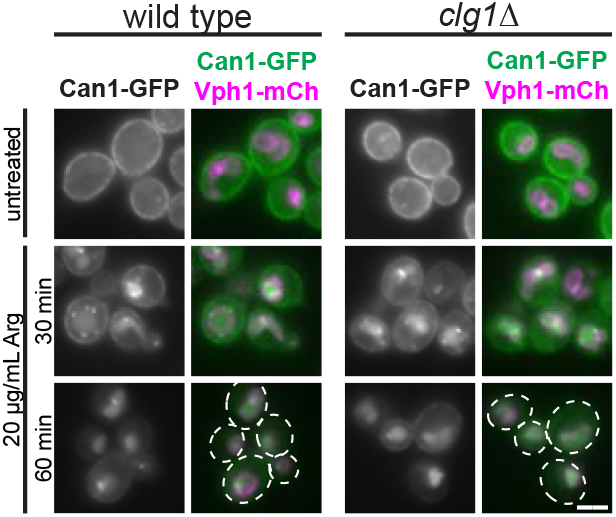
Mup1-GFP endocytosis in *clg1Δ.* Fluorescence microscopy of wild type or *clg1Δ* cells expressing Can1-GFP, with and without inducing endocytosis by treating with 20 μg/mL Arg for 30 and 60 min. Bar = 2 μm.

**Figure 5 – supplement 1.**
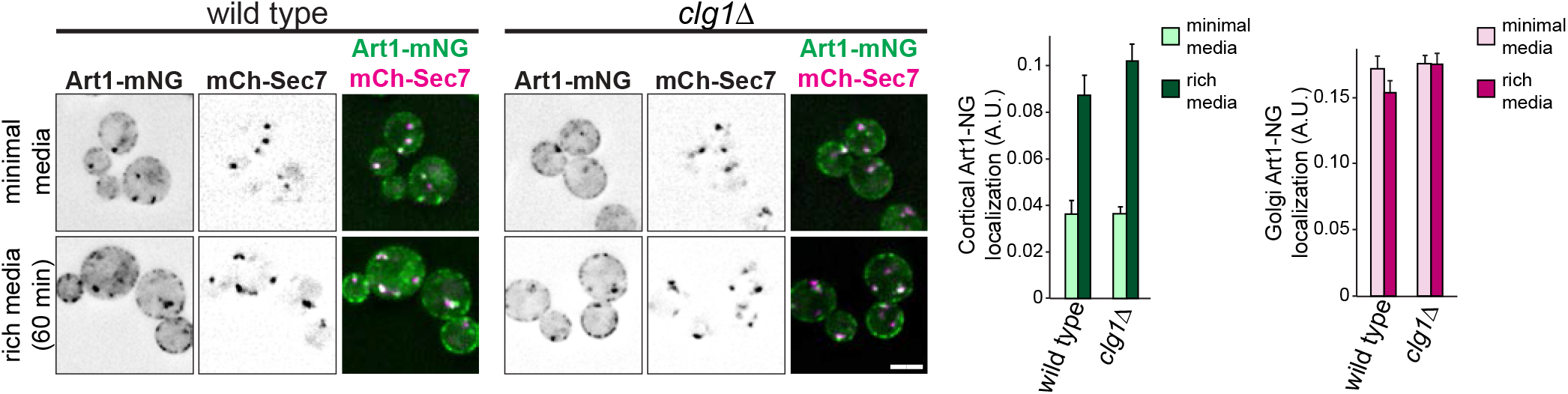
Art1 localization in *clg1Δ.* Localization of Art1-mNG in wild type and *clg1Δ* yeast in minimal media and after shifting to rich media for 60 min. Bar = 2 μm. (right) quantification of Art1-mNG PM and Golgi localization. Bars indicate 95% confidence intervals.

**Figure 6 – supplement 1.**
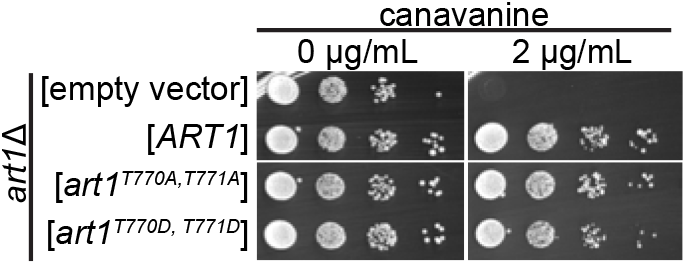
Art1 C-tail point mutant growth on canavanine. Serial dilutions of the *art1Δ* transformed with the indicated plasmids spotted on synthetic media containing the indicated concentration of canavanine and incubated at 26°C.

**Figure 6 – supplement 2.**
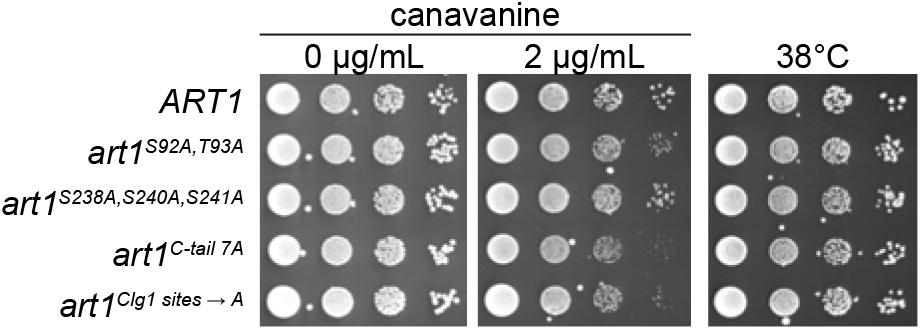
Art1 Ala point mutant growth on canavanine. Serial dilutions of the indicated strains spotted on synthetic media containing the indicated concentration of canavanine and incubated at 26°C or 38°C.

**Figure 6 – supplement 3.**
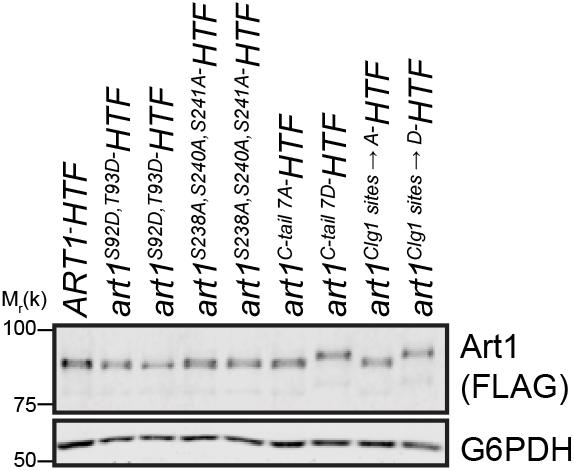
Art1 point mutant expression. Immunoblot analyzing the steady state expression of the indicated Art1 point mutants.

**Figure 6 – supplement 4.**
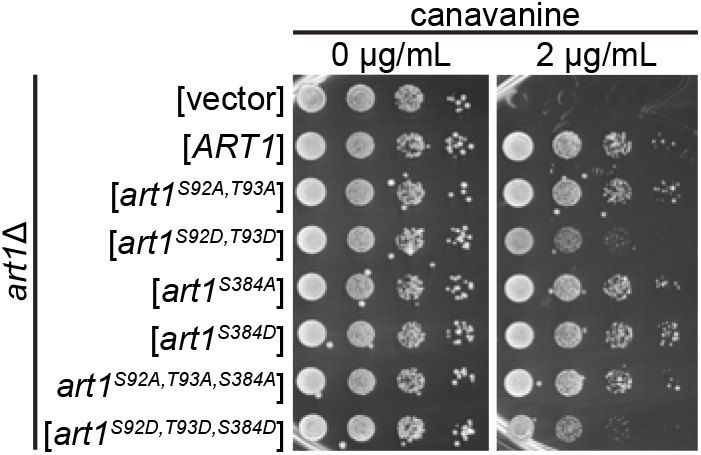
Art1 S384 point mutant growth on canavanine. Serial dilutions of the *art1Δ* transformed with the indicated plasmids spotted on synthetic media containing the indicated concentration of canavanine and incubated at 26°C.

**Figure 6 – supplement 5.**
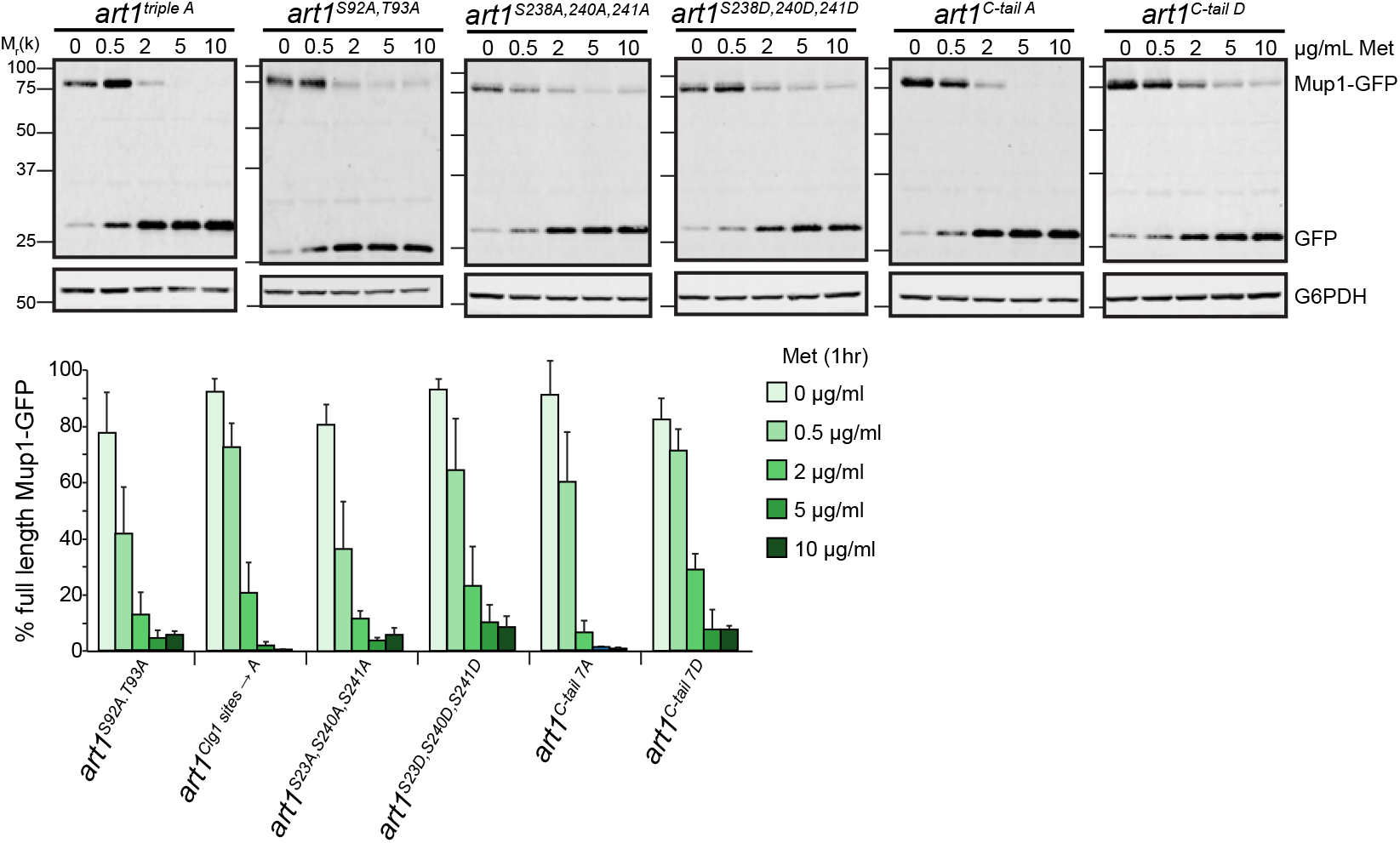
Mup1-GFP degradation in Art1 point mutants. (top) Immunoblot analysis of Mup1-GFP endocytosis induced with the indicated concentration of Met for 60 min in strains expressing the indicated Art1 mutant. (bottom) Band intensities from three biological replicates were quantified and expressed as the mean % full length Mup1-GFP remaining. Bars indicate 95% confidence intervals.

**Figure 6 – supplement 6.**
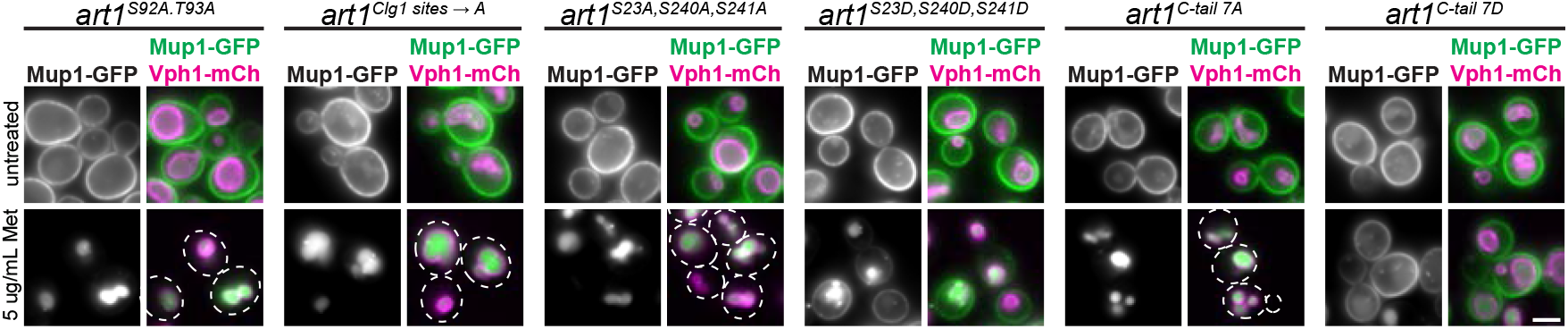
Mup1-GFP endocytosis in Art1 point mutants. Fluorescence microscopy of Mup1-GFP, with and without inducing endocytosis by treating with 5 μg/mL Met for 60 min in strains expressing the indicated Art1 mutant. Bar = 2 μm

**Figure 6 – supplement 7.**
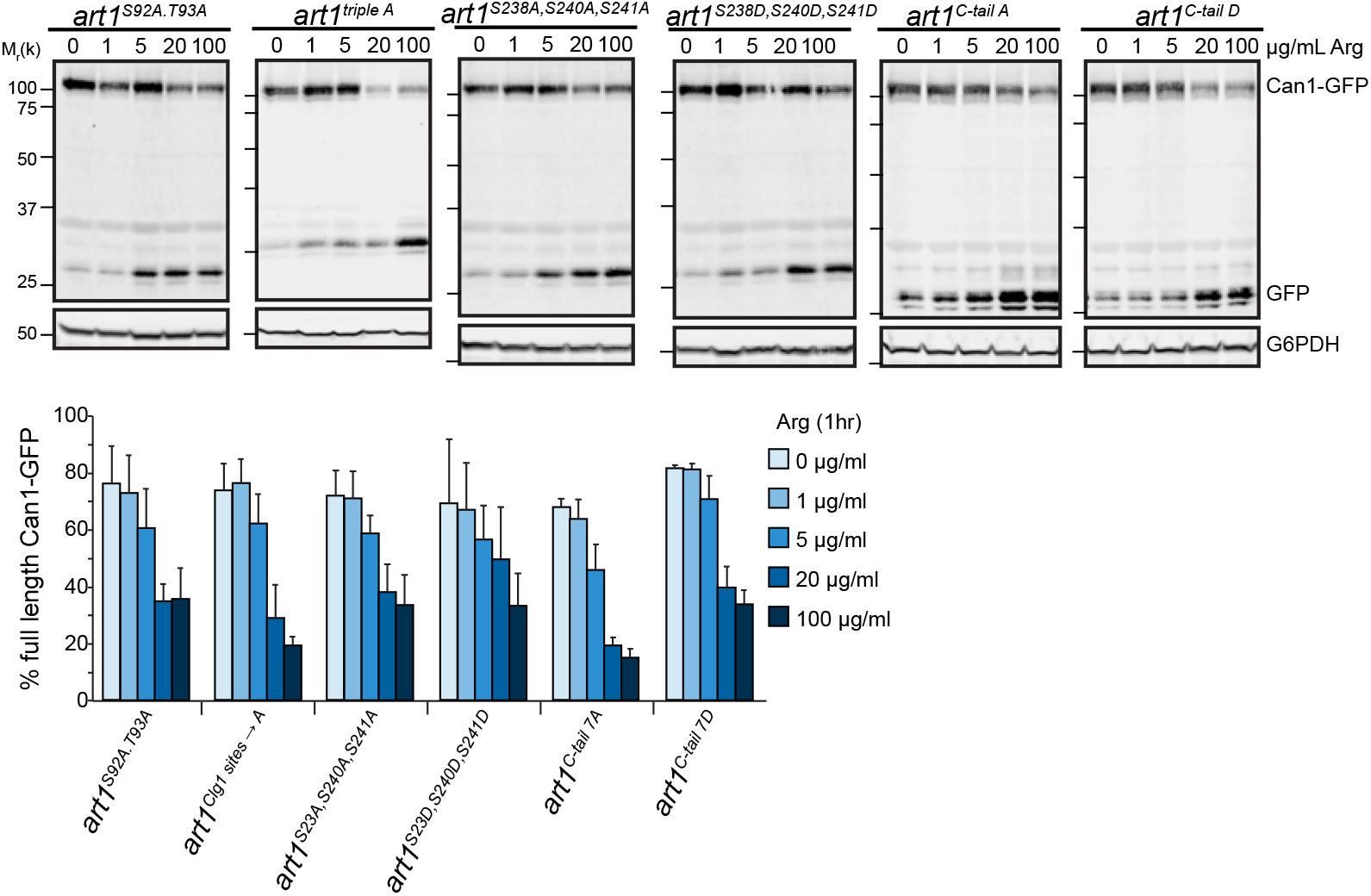
Can1-GFP degradation in Art1 point mutants. (top) Immunoblot analysis of Can1-GFP endocytosis induced with the indicated concentration of Arg for 60 min in strains expressing the indicated Art1 mutant. (bottom) Band intensities from three biological replicates were quantified and expressed as the mean % full length Can1-GFP remaining. Bars indicate 95% confidence intervals.

**Figure 6 – supplement 8.**
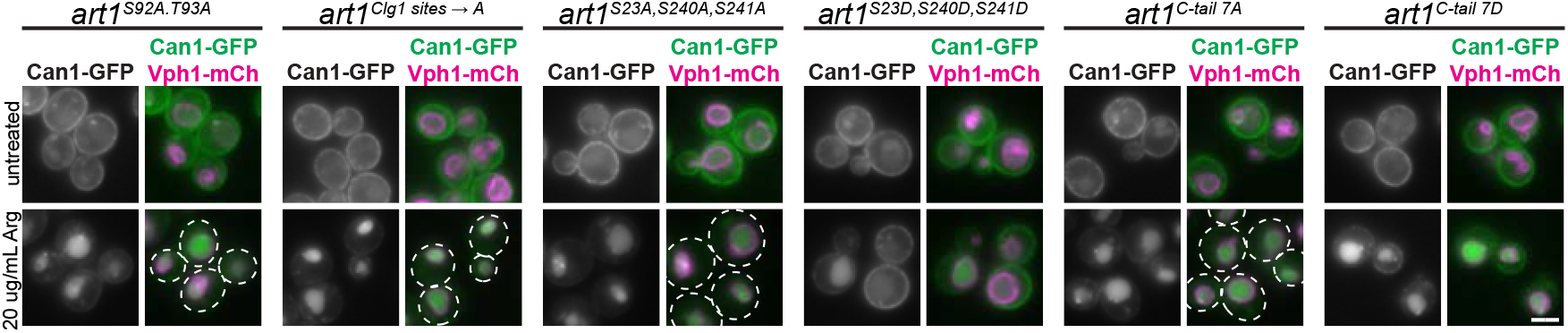
Can1-GFP endocytosis in Art1 point mutants. Fluorescence microscopy of Can1-GFP, with and without inducing endocytosis by treating with 20 μg/mL Arg for 60 min in strains expressing the indicated Art1 mutant. Bar = 2 μm

**Figure 7 – supplement 1.**
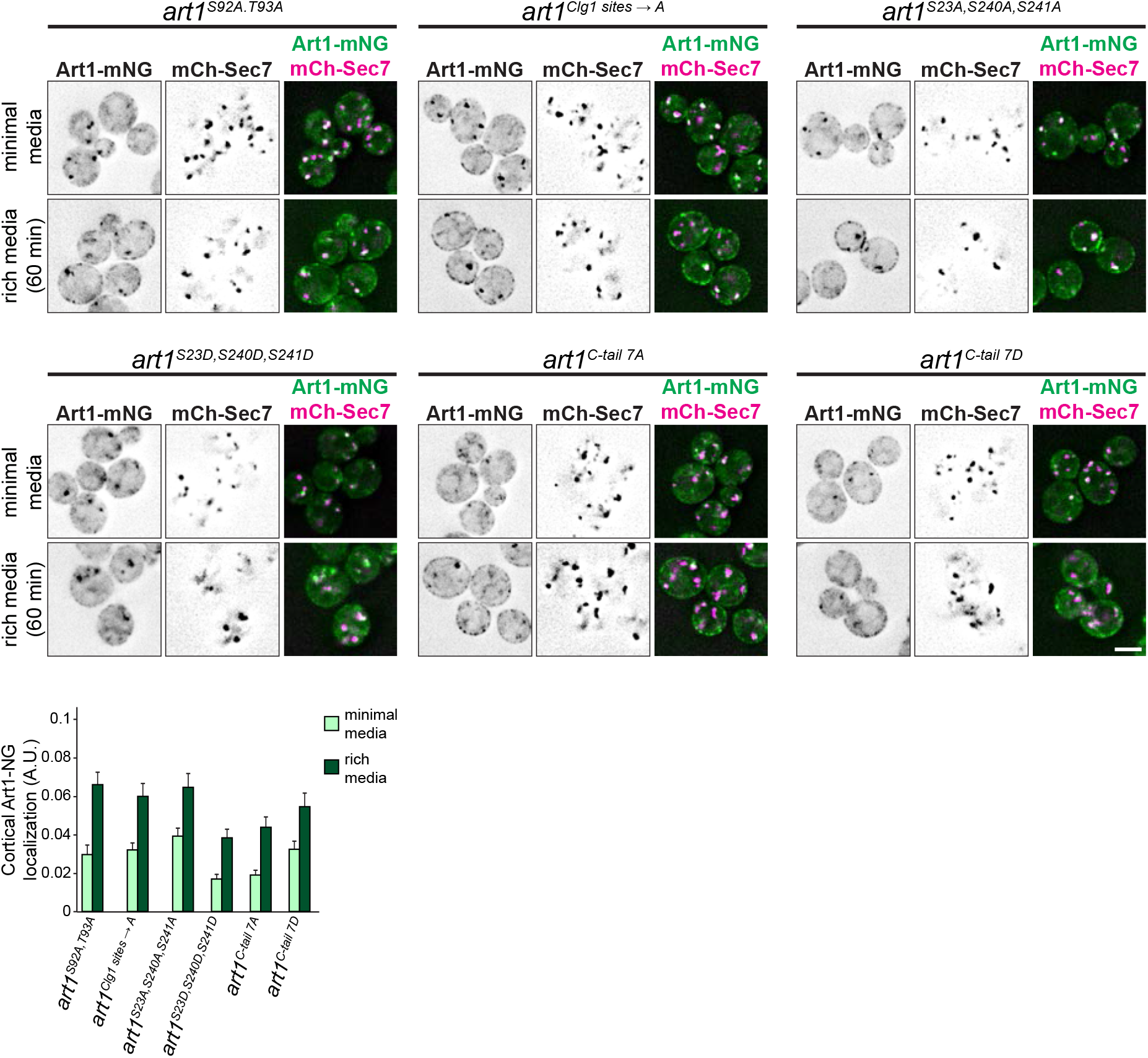
Art1 localization in *clg1Δ*. (top) Localization of Art1-mNG mutants in minimal media and after shifting to rich media. Bar = 2 μm. (bottom) quantification of Art1 cortical localization. Bars indicate 95% confidence intervals.

